# Eco-evolutionary dynamics of defense systems in mobile genetic elements: Cui bono?

**DOI:** 10.64898/2026.05.25.727639

**Authors:** Jaime Iranzo, Yuri I. Wolf, Eugene V. Koonin

## Abstract

**Background:** Mobile genetic elements (MGEs), including viruses, plasmids, and transposons, are major drivers of evolution in bacteria and archaea. Host-parasite conflicts drive the emergence of a broad variety of defense and counter-defense systems. Recent advances in metagenomics and functional annotation have shown that many defense systems are located on MGEs. The fact that MGEs are, essentially, genomic parasites raises an intriguing question: why do these parasites carry defense systems at high prevalence, often even higher than the host chromosome?

**Results:** We developed a simple mathematical model to investigate the factors that promote evolution of defense systems in MGEs and the ecological implications of MGE-encoded defense. Our analysis points to the strength of inter-MGE interference as a key determinant of the evolution of defense systems in MGEs. We identify two qualitatively distinct regimes, depending on the basic reproductive number in mixed coinfections. Weakly interfering MGEs tend to carry low-cost defense systems that enhance the survival of their hosts upon exposure to more damaging MGEs. Although these systems can be occasionally transferred to the host, they typically remain in MGEs. In contrast, strongly interfering MGEs, such as plasmids from the same incompatibility group, can carry high-cost defense systems that are detrimental to the host and the population as a whole, but help their carriers spread by actively replacing their competitors.

**Conclusions:** Analysis of our model shows that the key determinant of the evolution and spread of defense systems in MGEs is the strength of cross-MGE interference. Weakly interfering MGEs would serve as ‘MGE banks’, typically carrying low-cost defense systems that can benefit the host by protecting it from more damaging MGEs. In contrast, strongly interfering MGEs would carry costly defense systems that mediate inter-MGE conflicts but are deleterious to the host. These MGEs could serve as proving grounds for emerging defense systems, which might eventually become cost-effective once optimized by selection.

## Background

Evolution of all life forms, and prokaryotes in particular, involves the perennial co-evolution of organisms with mobile genetic elements (MGEs), such as viruses, transposons and plasmids [1-4]. This coevolution involves both the proverbial arms race and various forms of cooperation between MGEs and their hosts [4-6]. In the last few years, the development of dedicated computational and experimental approaches for the identification of defense systems, together with the advances in genomics and metagenomics, resulted in the discovery of an unexpected and unprecedented diversity of such systems in prokaryotes, with about 400 distinct ones identified, and counting [7, 8]. In the course of the host-parasite arms race, MGEs have evolved a commensurate diversity of counter-defense mechanisms, many of which remain to be characterized [9-11].

Notably, numerous defense systems were identified not only in bacterial and archaeal chromosomes but also in MGE genomes, especially, those of plasmids, large viruses, such as jumbo phages, as well as satellite viruses and proviruses [12-17]. The high prevalence of defense systems in MGEs, often higher than in non-mobile chromosomal regions, implies that defense systems are not simply occasionally hitchhiking on MGEs, but were exapted for specific roles beneficial for the latter. One of such roles can be counter-defense whereby MGE-encoded defense systems target and suppress host defenses. A case in point is the well-characterized type I-F CRISPR-Cas system encoded by Vibrio bacteriophage ICP1 and targeting the PLE elements that are involved in antiphage defense in these bacteria [18, 19]. Another, and possibly, more common role of MGE-encoded defense is inter-MGE conflicts. Examples include type IV and type V-M CRISPR-Cas systems whose primary role appears to be inter-plasmid competition [20-24]. In other cases, MGEs recruit only some components of a defense system and then, take advantage of the corresponding host-encoded defense machinery to redirect these against competitor MGEs. An example includes viral CRISPR mini-arrays that target competing viruses by hijacking host Cas proteins [25, 26]. Alternatively, MGEs can recruit defense systems for roles distinct from counter-defense such as RNA-guided transposition that is mediated by partially inactivated CRISPR-Cas systems carried by many Tn7-like and some Mu-like transposons [15, 27-31]. Sharing of defense systems by prokaryotic hosts and MGEs has been conceptualized as “guns for hire” whereby the same componentry is deployed by hosts for defense and by MGEs for counter-defense or other functions [32].

Most bacteria and archaea are hosts to multiple MGEs of different types that, together with the host, form a complex ecosystem with multiple interactions. Here, we ask: why are defense systems so often encoded by MGEs? Is it the host ‘outsourcing’ defense tasks or the MGEs themselves, launching defense systems against the host or other MGEs, that primarily benefit from the MGE-encoded defense systems? We systematically analyzed a simple eco-evolutionary model of a population of prokaryotes hosting competing MGEs to address these questions.

## Model

### General assumptions

We simulated a population of hosts and MGEs, with the dynamics on both sides described by a set of generalized replicator-mutator equations (see details below) [33]. In all variants of the model, we considered a well-mixed population that consists of MGE-free and MGE-carrier hosts. Free MGEs were not explicitly modeled. This is a natural choice for plasmids that do not exist in a cell-free state. In the case of viruses, the host-to-host transmission rate should be interpreted as an effective parameter that encompasses the rates of virus release and infection of new hosts.

The generalized replicator-mutator framework tracks the relative abundances of MGE-free and MGE-carrier hosts under the assumption that MGE transmission dynamics depend on relative abundances (as opposed to absolute abundances; Appendix A). Such dependency implies that MGE transmission rates are not limited by the total number of cells in the population, which is a reasonable approximation as long as the population density is moderate or high. Replicator-mutator equations admit stationary solutions in which the population grows, declines, or remains constant in size, making them a versatile tool to model population dynamics [34]. If the population size remains approximately constant (as it occurs in chemostats or in natural populations that have reached their saturation density for a given amount of resources), the distinction between relative and absolute abundances reduces to a scaling factor.

All variants of the model consider 2 classes of MGEs (henceforth A and B), characterized by their cost and transmission rates. Host chromosomes can harbor costly defense systems against one or both MGEs. Each MGE can also encode a defense system against the other MGE (Figure 1). Different variants of the model explore different options for the costs of MGEs and defense systems, including unequal costs for chromosome-encoded and MGE-encoded defense, and non-linear scaling of the costs with the number of MGEs and defense systems. The models also explore alternative scenarios of possible interference between MGEs cohabiting within the same host and the outcome of superinfection in which a host that already carries one MGE receives another MGE with a defense system that targets the resident MGE.

**Figure 1.**
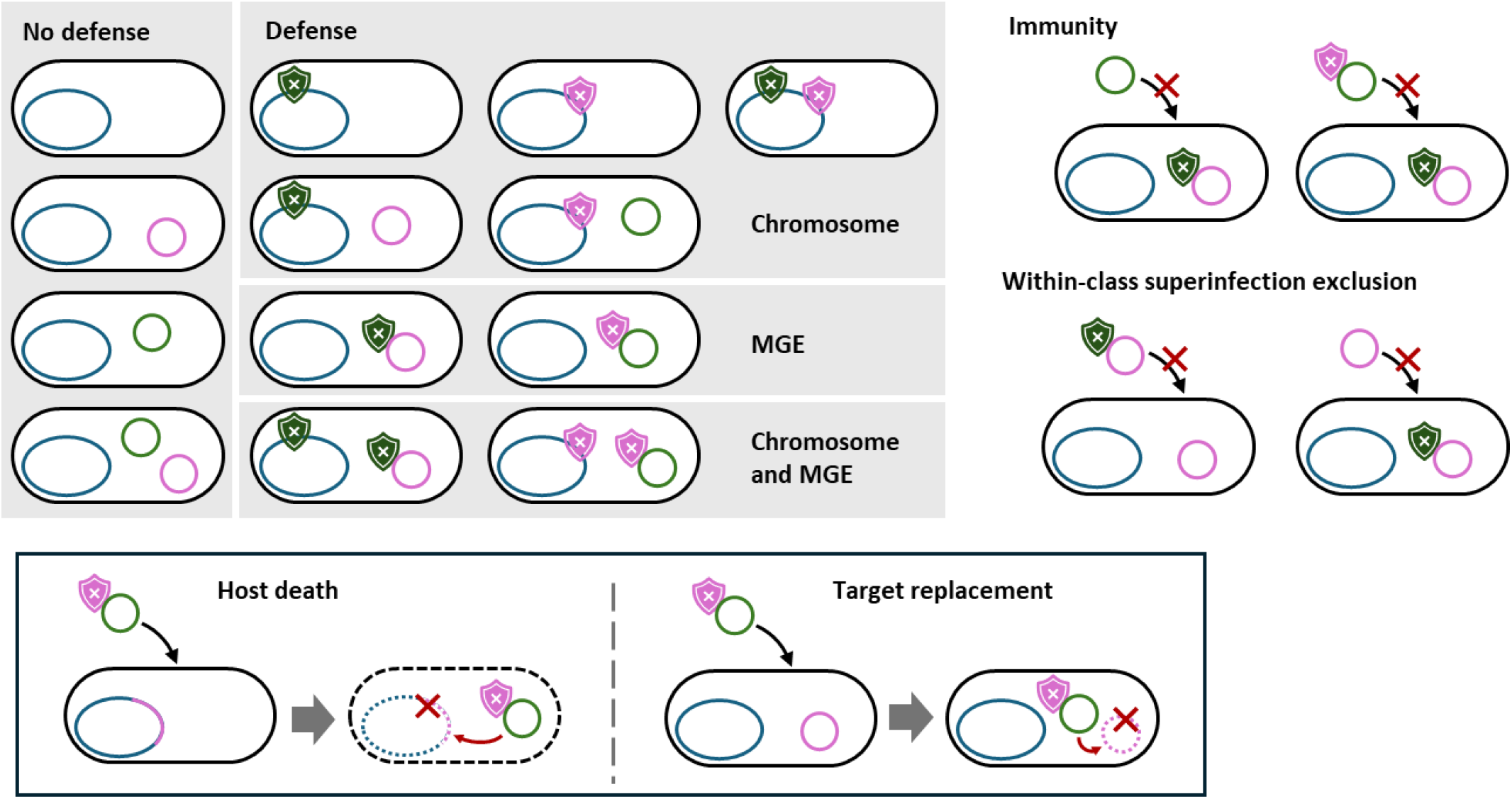
Schematic representation of the host and MGE classes and the infection processes simulated by the model.

### Generalized replicator-mutator equation for a population of hosts and MGEs

Let us consider a heterogeneous population of hosts, with *X*_*i*_ representing the number of hosts of a given class *i*. Considering that there are 2 classes of MGEs and that defense systems can be located on the host chromosome or on one or both of the MGEs, the total number of possible host classes is 13 (Fig 1 and Table 1).

**Table 1.** Host classes considered in the model. Subindices indicate the target of immunity, capital letters indicate which MGEs are present in a host. Numbers in parentheses indicate the index (*i*) assigned to each class in the model equations.

| MGE presence | Defense |  |  |  |
| --- | --- | --- | --- | --- |
|  | None | Chromosome-encoded | MGE-encoded | Chromosome and MGE |
| None | H (1) | H<sub>a</sub>(5), H<sub>b</sub> (6), H<sub>ab</sub> (7) | - | - |
| A | A (2) | H<sub>b</sub>A (8) | A<sub>b</sub> (10) | H<sub>b</sub>A<sub>b</sub> (12) |
| B | B (3) | H<sub>a</sub>B (9) | B<sub>a</sub> (11) | H<sub>a</sub>B<sub>a</sub> (13) |
| A and B | AB (4) | - | - | - |

The dynamics of the population can be generically represented with a system of 13 differential equations (one for each *X*_*i*_) of the following form:

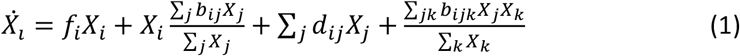

In these equations, 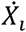 represents the total rate of change of host class *i* (in absolute abundance); *f*_*i*_ is the intrinsic replication rate of that class; *b*_*ij*_ represents the rate at which MGEs are transferred to or from hosts of other classes (*j*), respectively, increasing or decreasing the absolute abundance of class *i*; *d*_*ij*_ accounts for the production of hosts of class *i* directly from class *j* (for example, through the loss of an MGE or a defense system); and *b*_*ijk*_ represents the rate at which hosts of class *i* are produced via the transfer of MGEs between hosts of two different classes (*j* and *k* ≠ *i*).

Equation 1 can be expressed as a function of the relative abundances *x*_*i*_, where *x*_*i*_ = *X*_*i*_/ Σ *X* (Appendix A):

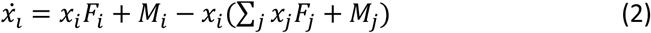

where *F*_*i*_ = *f*_*i*_ + Σ_*j*_ *b*_*ij*_*x*_*j*_ and *M*_*i*_ = Σ_*j*_ *x*_*j*_(*d*_*ij*_ + Σ_*k*_ *b*_*ijk*_*x*_*k*_). Equation 2 has the structure of a generalized replicator-mutator equation, in which *F*_*i*_ is a generalized fitness term and *M*_*i*_ is a generalized mutation term. In this formulation, the population dynamics remain invariant with respect to any additive fitness term that equally affects all host classes. This being the case, without loss of generality, we simply set the intrinsic replication rate in the absence of MGEs and defense systems to zero.

### Parameterization of the model

We assume that the cost of chromosome-encoded defense depends on how many MGEs are targeted. Thus, immunity against a single MGE has a cost *k*_1_ and immunity against both MGEs has a cost *k*_2_. This parameterization can account for all possible classes of epistasis among chromosome-encoded defense systems. If the cost of defense increases linearly with the number of targets (reflecting, for example, acquisition of independent defense systems), then *k*_2_ = 2*k*_1_. If defense against subsequent MGEs is less costly (for example, via acquisition of new spacers in a preexisting CRISPR array), then *k*_2_ < 2*k*_1_ (sublinear scaling). The extreme case *k*_2_ = *k*_1_ would correspond to a scenario in which immunity against the second MGE comes “for free” once the defense system protecting the host from the first MGE has been acquired, that is, the defense system provides full cross-protection. By contrast, if the cost of defense per target grows faster than linearly with the number of targeted MGEs, then *k*_2_ > 2*k*_1_ (superlinear scaling which could occur when defense systems overwhelm host resources).

The same considerations apply to the cost of MGEs. Let *c*_1_ be the average cost of a single MGE and *c*_2_ the joint cost of both MGEs. Then, for sublinear, linear, and superlinear scaling of the MGE cost, we have *c*_2_ < 2*c*_1_, *c*_2_ = 2*c*_1_, and *c*_2_ > 2*c*_1_, respectively. The model admits the possibility that the two MGEs have unequal fitness costs. We denote these by *c*_*A*_ (for MGE A) and *c*_*B*_ (for MGE B), with *c*_*A*_ ≤ *c*_*B*_. The average fitness cost of a single MGE becomes 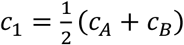 . For simplicity, we assume that MGE-encoded defense has an additive effect *k*_*p*_ on the total fitness cost, regardless of the presence of chromosome-encoded defense. The intrinsic growth rate of a host class is obtained by subtracting the cost of its MGEs and defense mechanisms. For example, for class H_a_B_a_, the growth rate is *f*_13_ = −*c*_*B*_ − *k*_1_ − *k*_*p*_.

MGE transfer is modeled through the interaction terms *b*_*ij*_ and *b*_*ijk*_. For simplicity, we assume that both MGEs have the same transfer rate, which may depend on the number of MGEs coinfecting the same host. The rate of transfer from hosts that contain a single MGE is *b* and the rate of transfer from coinfected hosts is *b*′ (per MGE). Thus, *b*′ < *b* corresponds to a scenario with coinfection interference. We assume superinfection exclusion within each class of MGE. In consequence, an MGE without defense systems cannot be replaced by a defense-encoding MGE of the same class (for example, a host carrying A cannot be superinfected by A_b_), and vice versa. Coinfection by the two MGE classes is only possible if none of them encodes defense against the other. Hosts carrying a defense system against an MGE cannot be infected by that MGE (complete immunity), regardless of whether the defense system is located on the chromosome or on a resident MGE.

A special situation occurs when a host that harbors one MGE and no defense is infected by an MGE of the second class, carrying a defense system that targets the resident MGE (for example a host carrying A could be infected by B_a_, which encodes a CRISPR array with a spacer against A). We considered two alternative scenarios to deal with these cases. In the first scenario, host death occurs, for example, because of the failure of DNA repair after defense-driven cleavage of chromosomally integrated MGEs, or because of inactivation of antitoxins encoded in the chromosome or in the resident MGE. In the second scenario, the resident MGE is removed or inactivated by the new one (defense-driven replacement). This scenario applies not only to non-integrative MGEs but also to integrative ones, provided that DNA repair mechanisms successfully maintain host genome integrity after cleavage of the MGE. A third possible scenario is that the resident MGE is not destroyed or silenced by the newly acquired defense system, but is simply tolerated. This would be the case of defense systems that only target modified DNA, such as type IV restriction-modification systems [35, 36]. This scenario can be assimilated into host death or target replacement, depending on the molecular details of interference among MGEs. For example, because plasmids of the same incompatibility group share replication and partitioning mechanisms, hosts coinfected by both MGEs will eventually lose one of them [37]. This outcome is analogous to the target replacement scenario, whereby the defense-encoding MGE superinfects and replaces the resident MGE in part of the offspring. Alternatively, interactions between some MGEs can exacerbate their fitness costs [38, 39], leading to the decline of coinfected host populations. In that case, superinfection by defense-encoding MGEs leads to host death.

We introduced a loss rate *d* that accounts for large deletions affecting multiple genes and leading to the loss of MGEs or chromosome-encoded defense systems. For simplicity, we did not consider cases in which MGEs lose their own defense systems but remain viable (although a realistic possibility, we assumed that this occurs at much lower rates). We initially set *d* = 0 and later explored the effect of introducing non-zero loss rates. Loss of MGEs and chromosome-encoded defense systems enters the replicator-mutation equations not only through the *d*_*ij*_ terms, but also as a negative contribution to the replication term *f*_*i*_ which is proportional to the number of MGEs and chromosome-encoded defense systems (Appendix B). We assumed that chromosome-encoded defense, once lost, cannot be regained (at least not at the time scales relevant to the model). This simplifying assumption greatly facilitates the interpretation of the results by making persistence of chromosome-encoded defense systems dependent on their fitness contribution (otherwise, defense could persist as a deleterious trait sustained by occasional regain).

Table 2 provides a summary of the model parameters and their interpretation. The explicit forms of the generalized replicator-mutator equations can be found in Appendix B.

**Table 2.** Parameters of the model.

| Symbol | Parameter |
| --- | --- |
| $b$ | Transfer rate per MGE (single-MGE infected hosts) |
| $b'$ | Transfer rate per MGE (coinfected hosts) |
| $c_A, c_B$ | Cost of MGE class A and B, respectively |
| $c_1$ | Average cost of a single MGE, equal to $(c_A + c_B)/2$ |
| $c_2$ | Cost of carrying both MGEs |
| $k_1$ | Cost of chromosome-encoded defense against a single MGE |
| $k_2$ | Cost of chromosome-encoded defense against both MGEs |
| $k_p$ | Cost of MGE-encoded defense |
| $d$ | MGE and defense loss rate |

## Results

### Population dynamics in the absence of MGE-encoded defense

To better comprehend the eco-evolutionary drivers and implications of MGE-encoded defense, we first explored simpler scenarios in which defense can only evolve in the host chromosome. As a starting point, let us consider a population with two classes of MGEs, A and B, and no defense systems. In this simple scenario, there are three possible stable regimes: (i) an MGE-free population, (ii) a population consisting of hosts carrying one class of MGE, and (iii) a population consisting of hosts carrying both MGEs (Appendix C). In the latter two cases, occasional MGE loss can lead to small fractions of MGE-free hosts and hosts with a single MGE, respectively, but those minority subpopulations are not self-sustaining. Which regime prevails, depends on the balance between the cost and transfer rate of each class of MGE.

To better illustrate this point, let us consider the MGE’s basic reproductive number (*R*_0_), defined as the average number of secondary infections produced by each infected host. In this model, the basic reproductive number is simply the ratio between the rate at which MGEs are transferred to new hosts (*b*) and the rate at which hosts infected by MGEs disappear from the population. The latter can occur through MGE loss (at a rate *d*) or selective displacement by MGE-less hosts (at a rate equal to the fitness cost of the MGE, *c* or *c*). Therefore, the basic reproductive number is 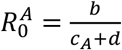 for MGEs of class A and 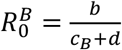 for MGEs of class B. Like in classical epidemiological models, MGEs cannot spread in the population unless their basic reproductive numbers are greater than one. Thus, if 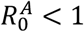 and 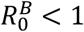, the host population will be free of MGEs. All parameter combinations explored in this work were chosen to ensure that 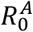 and 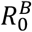 are both greater than 1 and, as a result, both MGE can spread if they are alone.

Similarly, we can define a basic reproductive number for mixed coinfections, that is, for hosts that contain both MGE classes. The expression in that case is 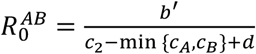 . In this expression, *b*^′^ is the rate at which coinfected hosts spread each of their MGEs, and c_2_ − min {*c*_*A*_, *c*_*B*_} is the additional fitness cost experienced by coinfected hosts compared to hosts with a single MGE. If MGEs do not interfere with each other’s transfer and fitness costs are additive (that is, *b*^′^ = *b* and *c*_2_ = *c*_*A*_ + *c*_*B*_), then, the basic reproductive number for the coinfection will be the same as for the pure infections 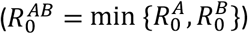 . As a result, if both MGEs can separately spread in the population, they will also be able to spread in the presence of each other and will coinfect the entire host population (note that 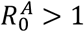 and 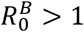 implies that 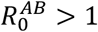). In contrast, if there is interference between the two MGEs, for example, due to reduced transfer rates (*b*^′^ < *b*) or increased fitness costs (*c*_2_ > *c*_*A*_ + *c*_*B*_), the basic reproductive number in coinfection will be lower than that in a pure infection 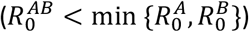 . If interference is so strong that 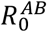 drops below 1 (while 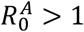 and 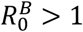), only one MGE will persist in the population. Typically, the MGE with the higher *R*_0_ will outcompete the other one although the opposite can happen if the initial composition is sufficiently skewed (see Appendix C.) Hereafter, we use “weak interference” and “strong interference” to refer to these two scenarios, characterized by 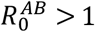 and 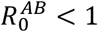 (while 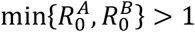), respectively. Weak interference includes the extreme case with no interference. In the absence of defense, weak interference leads to MGE coexistence whereas strong interference leads to competitive exclusion.

To provide biological context to these results, a paradigmatic case of strong interference among MGEs are plasmids of the same incompatibility group [37]. Other examples could include phage satellites that severely curtail replication of their helper phages [37], families of competing large plasmids, lytic phages, and, possibly, pseudolysogenic viruses that compete for limiting intracellular resources [16], as well as integrative elements that compete for the same insertion sites in the host chromosome [40]. Weak or no interference can be empirically inferred for pairs of MGEs that stably coexist in the same host. In particular, interactions between most lytic and lysogenic phages fall in the weak interference regime, given that the presence of prophages does not prevent replication of lytic phages except for most closely related ones. However, some prophages encode defense systems [41], in which case their interaction with susceptible lytic viruses falls under the strong interference regime.

Let us now suppose that some hosts encoding defense systems are introduced in the population. For simplicity, we assume that the defense systems are fully effective (that is, completely eliminate susceptible MGEs) and their mechanism of action does not involve growth arrest or cell death. Hosts carrying defense systems can either persist, not necessarily reaching fixation, or be outcompeted by susceptible hosts due to the cost of defense (Fig. 2A and S1). A necessary condition for the persistence of defense systems in the chromosome is that they are cost-effective, that is, the cost of defense (*k*_1_ or *k*_2_, for defense against one or both MGEs) must be lower than the cost of the MGE being targeted (*c*_*A*_, *c*_*B*_, or *c*_2_ for MGEs of class A, B, or both; see Appendix D for mathematical details). Thus, persistence of defense against a single MGE requires that *k*_1_ < *c*_*A*_ (dash-dotted blue lines in Fig. 2A) and persistence of defense against both MGEs in the same genome requires that *k*_2_ < *c*_2_ (dashed green lines in Fig. 2A). Depending on the relative costs of MGEs and defense systems, it can be less burdensome for the host to carry one MGE and a defense system against the other than to maintain defense against both MGEs (*k*_1_ < *c*_2_ − *c*_*A*_, dashed blue lines in Fig. 2A). Henceforth, we define 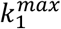 as the maximum cost-effective investment in chromosome-encoded defense.

**Figure 2.**
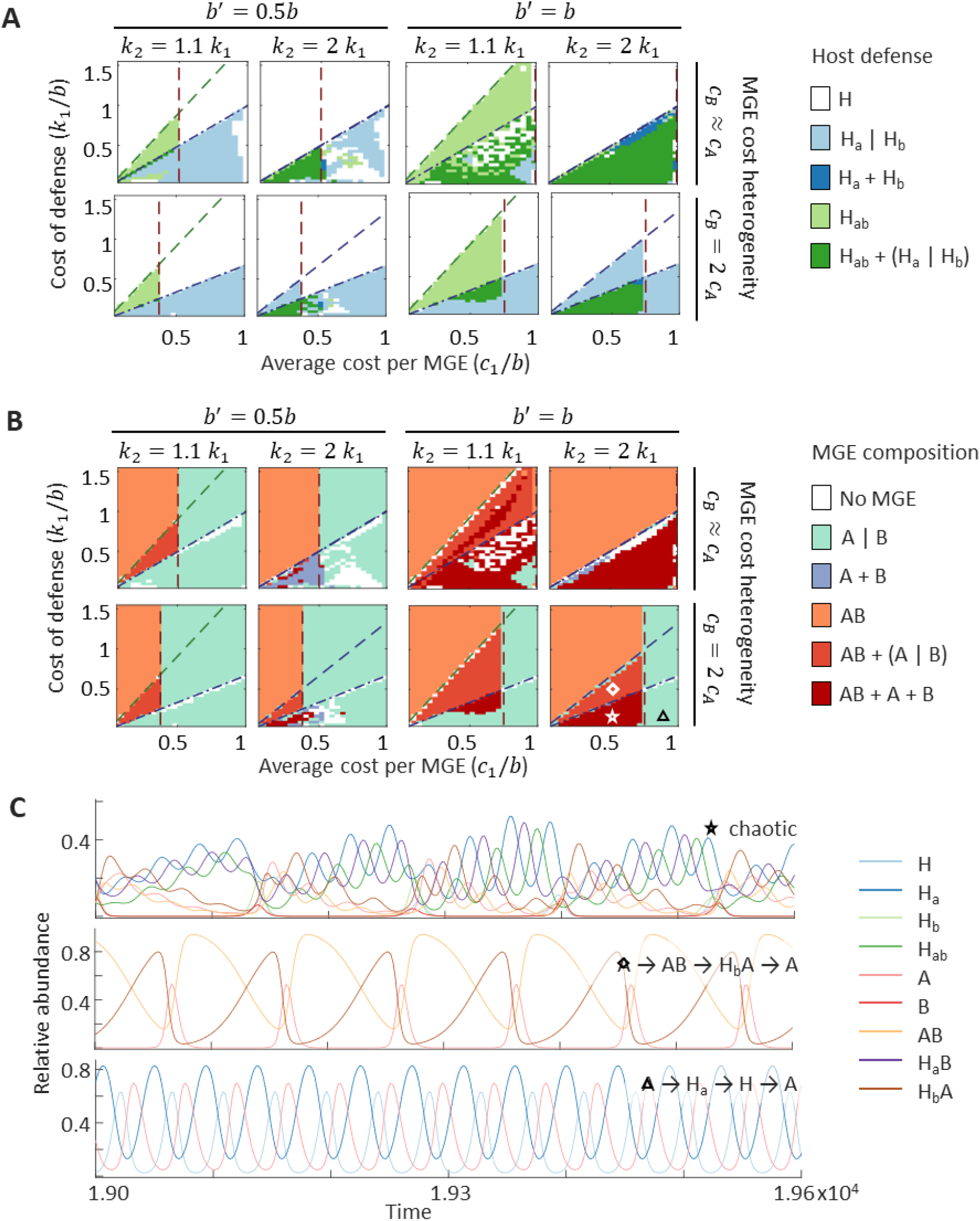
Evolution of chromosome-encoded defense and composition of the MGE population in the absence of MGE-encoded defense. A: Distribution of defense among host classes. “H_a_ | H_b_” denotes the presence of either H_a_ or H_b_. Brown dashed line: limit condition separating the weak (left) and strong (right) interference regimes, 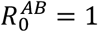. Dashed green line: limit condition for the spread of defense against both MGEs, *k*_2_ = *c*_2_. Dash-dotted blue line: limit condition for the spread of defense against a single MGE, *k*_1_ = *c*_*A*_. Dashed blue line: limit condition for the spread of defense against one MGE in hosts already carrying the other MGE, *k*_1_ = *c*_2_ − *c*_*A*_. B: Composition of the MGE population. “A | B” denotes the presence or MGE classes A or B, but not both. Reference lines as in A. C: Three representative examples of cyclical population dynamics in the absence of MGE-encoded defense. The parameter combinations that produce each time series are indicated by a star, a diamond, and a triangle in B. All simulations were run with zero loss rate (*d* = 0).

Chromosome-encoded defense can lead to diverse population dynamics (Fig. 2B-C and S2). Besides homogeneous stationary states, for many parameter combinations, the population displays cyclic and complex oscillations involving alternating subpopulations of immune and susceptible hosts (Fig. 2C). Oscillations are driven by the delayed accumulation and loss of defense systems following the spread and decline of MGEs. The simplest cycles involve three classes of hosts: susceptible hosts are first invaded by MGEs and then outcompeted by immune hosts, causing the decline of the MGEs and the subsequent recovery of the susceptible host population. More complex cycles can involve both classes of MGEs, as well as single and doble immunity, sometimes oscillating in non-periodic, seemingly, chaotic orbits. In the absence of gene loss, oscillatory dynamics are almost universal. Otherwise, oscillatory dynamics are still widespread in the strong interference regime (Fig. S3). In some regions of the parameter space, the final composition of the population is highly sensitive to the initial conditions. Chromosome-encoded defense does not generally lead to MGE extinction although there are some exceptions associated with a collapse of the oscillatory dynamics (white regions in Fig. 2B).

### Evolution of MGE-encoded defense systems

Like in the case of chromosome-encoded defense, evolution of MGE-encoded defense is cost-dependent. To facilitate the interpretation of the results, we will focus on the maximum cost 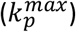 that is compatible with the persistence of MGE-encoded defense. To justify this choice, one can think of an arms race scenario in which co-evolution of MGE-encoded defense and anti-defense mechanisms leads to ever growing investment in defense. The value of 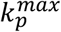 would then represent the maximum investment that can be reached before defense becomes inefficient.

The values of 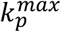 and 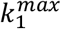 impose an upper bound on the cost of defense systems encoded in MGEs and in the chromosome, respectively. Simulations show that these two values do not coincide in general and sometimes even follow opposite trends. In the absence of chromosome-encoded defense, 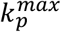 qualitatively depends on the strength of interference between the two MGEs (Fig. 3A and S4). In the weak interference regime, 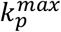 linearly increases with the cost of MGEs and never surpasses the maximum cost of chromosome-encoded defense (that is, 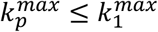). As a result, defense mechanisms that enter a population via MGEs in this regime will remain cost-effective if transferred to the chromosome. In the strong interference regime, spread of MGE-encoded defense is only expected under the target replacement scenario, that is, if defense systems can help the invading MGE eliminate the resident MGE (see *Model* and Fig. 1 for further details). In that case, defense systems located on MGEs can evolve at substantially higher costs than those located on the chromosome. The picture becomes more complicated if the host also harbors defense systems on the chromosome because the interplay between the two sources of defense makes 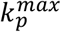 also dependent on the cost of chromosome-encoded defense (Fig. 3B and S5). However, despite some quantitative differences, the same general trends persist: MGE-encoded defense of similar or lower cost (that is, similar or higher efficiency) than chromosome-encoded defense can spread under weak interference conditions, especially in MGEs with unequal costs (in that case, defense is typically encoded by the least costly MGE and targets the most burdensome MGE). In contrast, ultra-costly (cost-ineffective) defense can spread among strongly interfering or incompatible MGEs only under the target replacement scenario. These trends are robust with respect to introducing nonzero loss rates in the model (Fig. S6-7).

**Figure 3.**
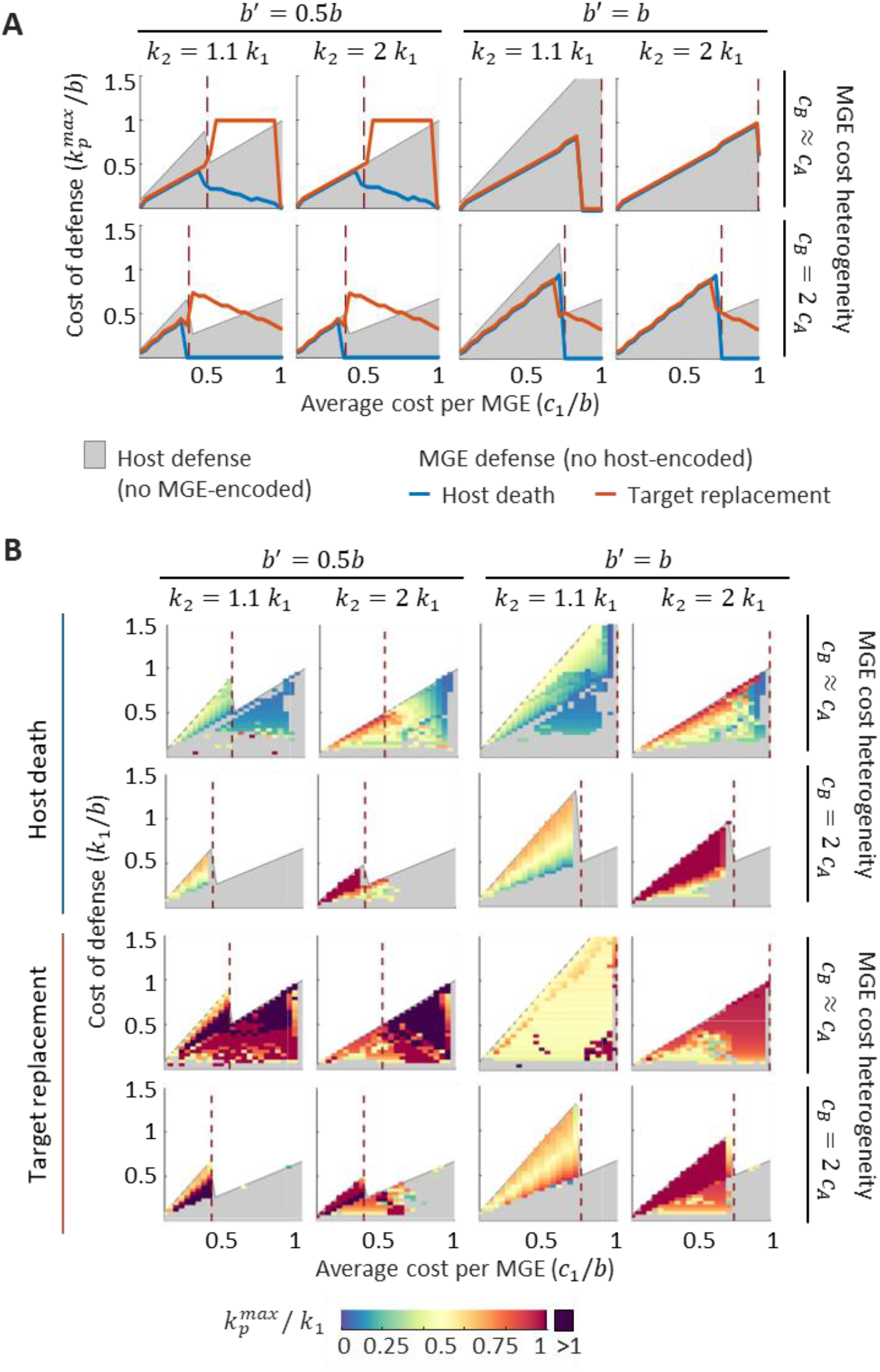
Maximum cost of defense 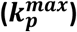 compatible with the spread of MGE-encoded defense, in the absence (A) or presence (B) of chromosome-encoded defense. Parameter regions where chromosome-encoded defense would be cost-effective are shown in gray. In B, colors indicate the ratio 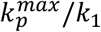to account for the fact that the maximum cost of MGE-encoded defense in the presence of chromosome-encoded defense depends on the cost of the latter. Brown dashed line: limit condition separating the weak (left) and strong (right) interference regimes, 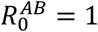. All simulations were run with zero loss rate (*d* = 0).

The distinct outcomes of defense evolution predicted under weak and strong interference can be largely explained by differences in the relative contributions of vertical and horizontal transmission to the spread of MGE-encoded defense. In the weak interference regime, all susceptible hosts are initially coinfected by non-defense-encoding variants of both MGEs. Defense-encoding MGEs cannot be horizontally transferred to these hosts because of within-class superinfection exclusion. As a result, they can only spread via vertical transmission and host-level selection, which will occur only if the MGE-encoded defense provides a fitness benefit to the host (with respect both to the cost of chromosome-encoded defense and the other MGE). Thus, the spread of MGE-encoded defense requires that *k*_*p*_ ≤ *k*_1_, *k*_*p*_ ≤ *k*_2_ − *c*_*A*_, and *k*_*p*_ ≤ *c*_2_ − *c*_*A*_, or in a more compact form, *k*_*p*_ ≤ min{*k*_1_; *k*_2_ − *c*_*A*_; *c*_2_ − *c*_*A*_} (we omitted the effect of gene loss for simplicity; see Appendix E for a formal derivation). Evolution of defense systems in weakly interfering MGEs always aligns with the interests of the host because the entire course of evolution is defined by host-level selection. In contrast, if interference is strong, the spread of MGE-encoded defense occurs through a combination of host-level selection and horizontal transfer between hosts containing a single MGE. Superinfection scenarios are critical in this regime because they determine the potential of defense-encoding MGEs to invade hosts that already contain the opponent MGE (typically, the least costly one), resulting in increased relative contribution of horizontal versus vertical transmission. Under the conditions of strong interference and target replacement, defense-encoding MGEs of class B (B_a_ in the notation of Fig. 1) can invade an all-A population if their horizontal transfer rate (*b*) is greater than the difference between its total cost (*c*_*B*_ + *k*_*p*_) and the cost of the resident MGE (*c*_*A*_), leading to the condition *k*_*p*_ ≤ *b* + *c*_*A*_ − *c*_*B*_. By rewriting this expression as 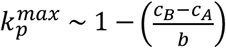, it becomes clear that ultra-costly defense in the target replacement scenario is more likely to evolve among MGEs that have similar (and/or large) *R*_0_. In contrast, under the host death scenario, horizontal transfer does not contribute to the spread of MGE-encoded defense. As a consequence, evolution of MGE-encoded defense becomes unfeasible (it would require that *k*_*p*_ ≤ *c*_*A*_ − *c*_*B*_, which cannot be satisfied as long as *c*_*A*_ < *c*_*B*_). By formalizing and extending these arguments, in Appendix E, we obtained analytical expressions that match the values of 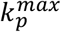 in Fig. 3A.

### Long-term coevolution and transfer of MGE-encoded defense to the chromosome

So far, we have been treating chromosome-encoded and MGE-encoded defense independently. However, defense systems originally carried by MGEs can be integrated into the host chromosome and domesticated. Likewise, host defense systems can be co-opted by MGEs. Considering such transfers, what would be the outcome(s) of co-evolution in the long term? Will MGE-borne and chromosome-borne variants of the same defense system persist, or will selection promote one over the other?

To address these questions, we investigated the fate of anti-MGE defense systems upon transfer from MGEs to the chromosome or vice versa. Within the model, two defense systems (MGE-encoded and chromosome-encoded ones) with the same cost and the same target are indistinguishable. Thus, the dynamics of recently transferred systems can be simulated by setting *k*_*p*_ = *k*_1_. To facilitate the interpretation of the results, here we focus on 4 representative cases that qualitatively cover most of the observed outcomes (Fig. 4); a comprehensive analysis is presented in Fig. S8-9.

**Figure 4.**
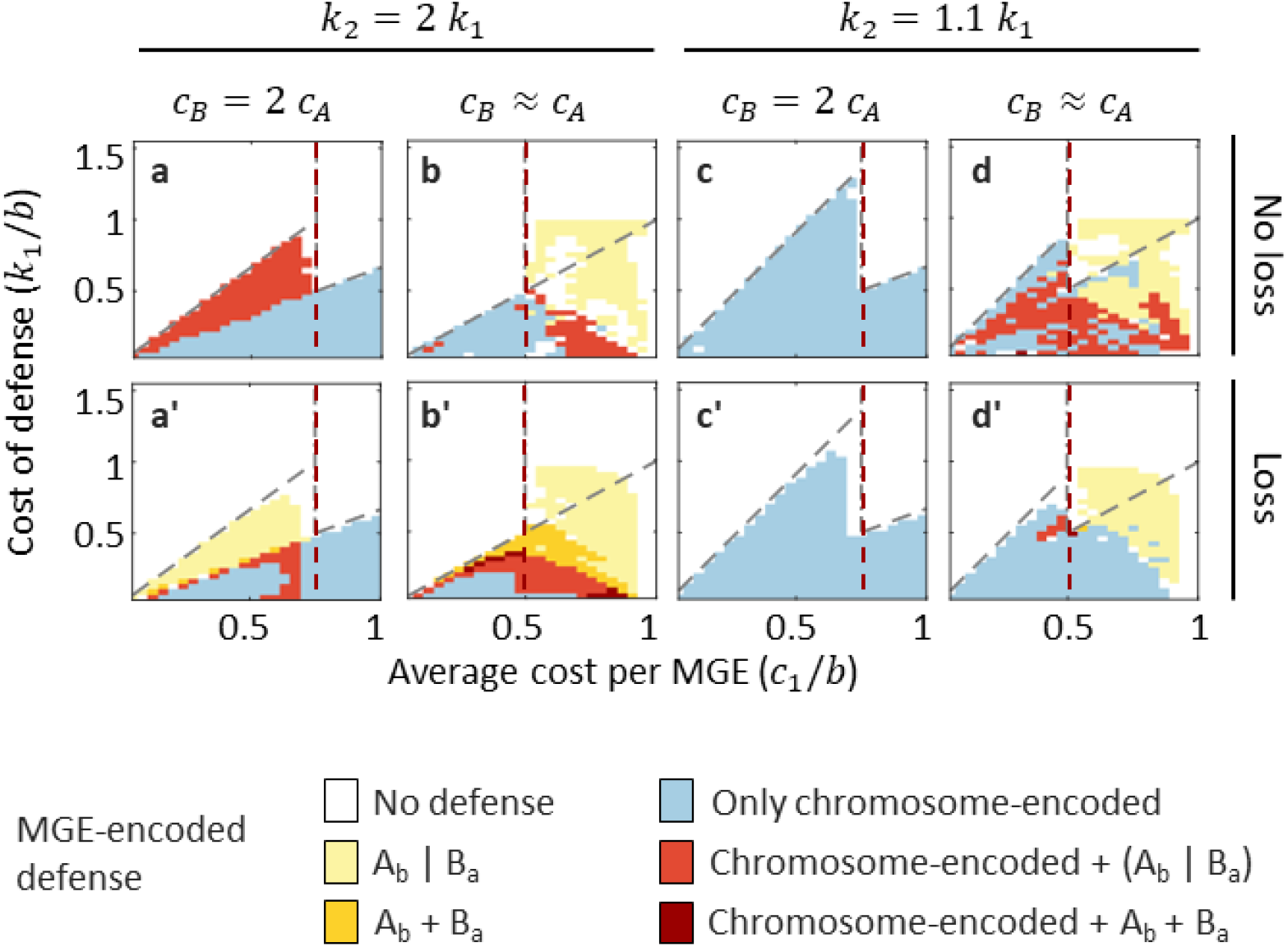
Expected location of defense in the long-term, considering the possible transfer of MGE-encoded defense systems to the chromosome and vice versa. “A_b_ | B_a_” indicates that only one of the two MGEs will encode a defense system. Brown dashed line: limit condition separating the weak (left) and strong (right) interference regimes, 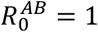. Dashed lines separate regions in which chromosome-encoded defense is cost-effective or not (below and above the line, respectively). Simulations were run with deletion rate *d* = 0 (no loss) and *d* = 0.01 (loss), and the same cost for MGE-encoded defense and chromosome-encoded defense (*k*_*p*_ = *k*_1_).

Plots a-a’ and b-b’ in Fig. 4 illustrate the case where the cost of defense linearly increases with the number of targets. In practice, such (approximately) linear scaling can be expected if defense systems targeting each MGE act independently of each other. The first plot (a) corresponds to two weakly interfering MGEs with unequal costs, such as a benign plasmid or prophage and a virulent phage. In the absence of gene loss, and for a relatively broad region of the parameter space, defense systems targeting the costliest MGE can persist in the benign MGE or in the chromosome. However, chromosome-encoded defense is lost if a nonzero deletion rate is introduced in the model (a’). This loss of defense systems results from the additional cost experienced by hosts that already harbor a defense system and are infected by an MGE that encodes the same defense system. The asymmetry between chromosome-borne and MGE-borne defense ultimately emerges from the different timescales for MGE reinfection and regain of chromosome-encoded defense after gene loss. Because the latter occurs at much slower rates, losing chromosome-encoded defense effectively reduces the burden of redundancy in the long term, whereas losing MGE-encoded defense does not (redundancy would reappear as soon as defense-encoding MGEs reinfects the host). All in all, despite transfer of defense systems from MGEs to the chromosome being feasible, the transferred copies are counter-selected, so that MGEs serve as the long-term reservoir of defense. Given that the host directly benefits from those defense systems, this scenario can be interpreted as a case of defense ‘outsourcing’, a case of a mutualistic relationship between the host and the MGE. The second set of plots (b-b’) corresponds to two strongly interfering MGEs with similar costs, such as two plasmids from the same incompatibility group (a qualitatively similar outcome is also observed for strongly interfering MGEs with unequal costs, see Fig. S8-9). In this scenario, defense is most often restricted to the MGEs, even in some cases where it would also be cost-effective for the host (yellow region under the dashed gray line). Only if sufficiently low-cost, defense systems evolving in this context can be transferred to the chromosome and would stably coexist with MGE-borne variants.

Plots c-c’ and d-d’ apply to scenarios in which defense against the second MGE comes at a lower cost once the host has defense against one MGE. CRISPR-based immunity obtained through acquisition of spacers in a preexistent CRISPR array would fall in this category. As shown in plots c-c’, long-term maintenance of this type of defense in weakly interfering MGEs is less likely although it can occur if they have substantially unequal costs (in that case, defense can be stably maintained both in the chromosome and the MGE, Fig. S8-9). In the case of incompatible or strongly interfering MGEs (d-d’), the situation is qualitatively similar to that obtained with additive fitness cost although, in this case, MGE-encoded defense systems are more likely to be completely transferred to the chromosome.

An alternative perspective on these results is to analyze the context in which a defense system of a given cost can be found. Thus, ultra-costly (or cost-ineffective) defense systems will only be found in incompatible MGEs, which use them as weapons to outcompete and displace each other. Cost-effective innate immunity systems will be most often found in benign MGEs targeting more virulent MGEs, although they can also appear in incompatible MGEs targeting each other. Although these defense systems can be transferred to the chromosome, their prevalence will generally be higher in MGEs, which serve as a reservoir. Finally, cost-effective adaptive immunity systems, that is, CRISPR-Cas, will generally occur in the chromosome or in strongly interfering MGEs.

### Population-level effects of MGE-encoded defense

MGE-encoded defense evolves in the context of inter-MGE conflicts. It is known that such conflicts can lead to traits that are not necessarily beneficial neither to the parties involved nor the host, a situation that is analogous to the Prisoner’s Dilemma in game theory [42-46]. In this context, we investigated the population-level effects of MGE-encoded defense by comparing the prevalence of each MGE and the growth rate of the host when the only source of defense is chromosome-encoded (by setting A_b_ = B_a_ = 0 in the simulations) and when MGE-encoded defense is also available. As in the previous section, we set *k*_*p*_ = *k*_1_ to account for the possibility that defense systems originally introduced in the population through MGEs are transferred to the chromosome and vice versa.

The effect of MGE-encoded defense on population composition fully depends on the extent of interference among MGEs (Fig. 5 top and S10). In the weak interference regime (left of the vertical red dashed lines in Fig. 5), MGE-encoded defense has no detectable effect on population composition. In contrast, under strong interference, MGE-encoded defense promotes diversity by fostering coexistence of MGEs. Such coexistence occurs at the population level, with different MGEs residing in different host cells. Population-level coexistence of MGEs can involve either cyclic replacement dynamics or emergence of stable subpopulations. Although coexistence sometimes involves reciprocal targeting, it is not essential for coexistence (Fig S11).

**Figure 5.**
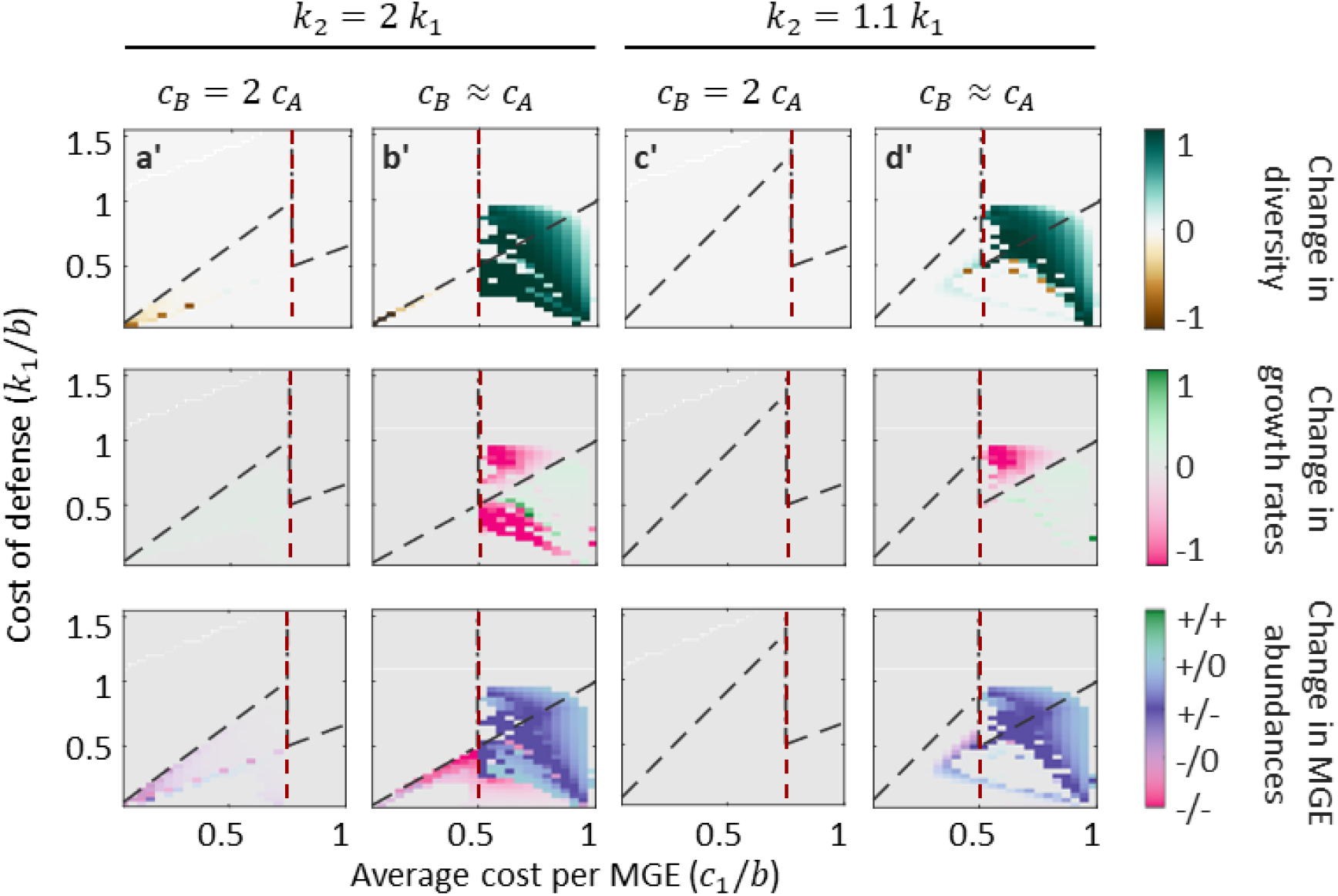
Effect of MGE-encoded defense on MGE diversity (top), overall fitness (middle), and local MGE fitness (bottom). Changes in diversity were quantified by comparing the Shannon index in the steady state of simulations with and without MGE-encoded defense. Overall fitness differences were quantified as the change in the logarithm of the growth rate with and without MGE-encoded defense. Changes in the local fitness of MGEs were quantified by calculating the differences in their relative abundances after introducing MGE-encoded defense. To obtain the Shannon index and the relative abundances, all variants (defense-free and defense-encoding) of each class of MGE were jointly considered. Vertical dash-dotted line: limit condition separating the weak (left) and strong (right) interference regimes, 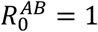. Dashed lines separate regions in which chromosome-encoded defense is cost-effective or not (below and above the line, respectively). All simulations were run with nonzero deletion rate (*d* = 0.01) and same cost for MGE-encoded defense and chromosome-encoded defense (*k*_*p*_ = *k*_1_).

From the perspective of the host, MGE-encoded defense is generally neutral or deleterious (Fig. 5 middle and S12). MGE-encoded defense is neutral when it evolves in the context of weakly interfering MGEs or under the host death scenario. Such neutrality is directly connected to the way in which host-level selection drives the spread of defense-encoding MGEs in these conditions (to spread via host-level selection, defense cannot be deleterious; neutrality ensues when the investment in defense reaches its maximum possible value 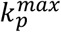). The deleterious effects of MGE-encoded defense are always associated with strong interference As explained above, in the strong interference regime (in the absence of defense), only one class of MGEs can persist in the population. The worst-case scenario for the host occurs when the MGEs that persist belong to the costliest class, B, whose cost is *c*_*B*_. Starting from that composition, MGEs of class A can invade the population via HGT if they encode defense against class B. The cost of these MGEs is *c*_*A*_ + *k*_*p*_, with 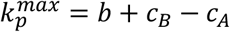 (Appendix E). Therefore, the replacement of one class of MGEs by the other will result in a net increase in the cost for the host, from *c*_*B*_ to *c*_*B*_ + *b*.

Thus, although MGE-encoded defense helps hosts remove the costliest MGE from the population, the total cost is greater than the benefit. As discussed above, such defense systems are ultra-costly or cost-inefficient for the host. Finally, although there exist small parameter regions in which MGE-encoded defense provides a net growth benefit over chromosome-encoded defense, this situation is rare and is restricted to highly specific scenarios, representing less than 0.2% of all the parameter combinations (Fig. S12).

From the perspective of the MGEs, defense systems display a broad range of fitness effects (Fig. 5 bottom and S13). Asymmetric outcomes in which only one MGE receives a benefit (that is, one MGE becomes more prevalent while the other one declines or remains unchanged) are typical in the strong interference regime. In contrast, in the weak interference regime, MGE-encoded defense most often leads to ‘no winner’ scenarios (including neutral, neutral-loss, or loss-loss outcomes).

Prevalence provides a quantitative measurement of local fitness, which can be understood in terms of the performance of each MGE within the population. However, from a global perspective, emergence of new MGEs that could spread beyond the local population and colonize new hosts depends on the overall growth rate. As discussed above, the effect of MGE-encoded defense on the host growth rate is generally neutral or negative, reducing the overall yield and, with it, the potential for dispersal of defense-encoding MGEs to other populations. This apparent paradox underscores the selfish nature of MGE-encoded defense, which contributes to competition among MGEs in local populations but, at a global (metapopulation) scale, becomes detrimental for their carriers.

## Discussion

The carriage of defense systems by MGEs creates a complex ecosystem in which the interests of the hosts and the MGEs can align or collide, depending on the system parameters. Analysis of our model points to the strength of cross-MGE interference as a key determinant of the evolution and spread of defense systems in MGEs. We identified two qualitatively distinct regimes, depending on the basic reproductive number associated with mixed coinfections 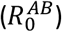 . Weakly interfering MGEs can propagate together in susceptible host populations 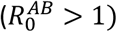 . These MGEs, such as compatible plasmids and temperate phages, can carry defense systems that enhance the survival of their hosts upon exposure to more damaging (but not necessarily incompatible) MGEs. Therefore, evolution of MGE-encoded defense in the weak interference regime generally aligns with the interests of the host. In contrast, strongly interfering MGEs cannot propagate together (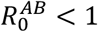 despite 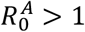 and 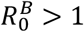) and their dynamics are driven by competitive exclusion. Those MGEs, for example, plasmids from the same incompatibility group, can carry ultra-costly (or cost-inefficient) defense systems that help them actively replace competitors. To accomplish such replacement, ultra-costly defense systems must be able to not only prevent the propagation of the competing MGE when it arrives in a cell carrying the defense-encoding MGE, but also destroy or silence a resident competing MGE without inflicting too much damage on the host.

MGE-encoded defense systems that evolve in the weak interference regime are equivalent to those on the chromosome, in the sense that they are subject to the same selective pressures and cost-efficiency conditions. Because of that, these systems could be readily “domesticated” (through integration followed by degeneration of the mobility-encoding regions of the MGE) and become part of the chromosome. Domestication is advantageous to the host because it allows keeping the defense system while dispensing of the cost of the MGE. Despite that potential benefit, analysis of our model suggests that complete transfer of MGE-encoded defense to the chromosome is not always evolutionarily stable. Indeed, if defense-encoding MGEs are sufficiently prevalent and benign, domestication could cause the hosts to carry redundant copies of defense systems, those that have been domesticated and those brought about via reinfection by the original MGE. To prevent such redundancy, defense outsourcing to MGEs could be a more efficient strategy than domestication. Furthermore, such outsourcing has a population-level effect, facilitating dissemination of defense systems across the host population. This phenomenon could, at least, in part explain why RM and other classes of defense systems are often more prevalent in plasmids, integrative elements and prophages than in regions of host chromosomes unrelated to MGEs. Defense outsourcing would result in a lower-than-expected redundancy between MGE-encoded and chromosome-encoded defense repertoires, a prediction that is testable by single-cell genomics approaches.

Type IV and subtype V-M CRISPR-Cas systems, that are typically found in plasmids and some viral genomes, differ from the great majority of CRISPR-Cas systems in their mechanism of action. Most of these MGE-associated variants do not cleave the target but only bind to the cognate sequence, blocking transcription or replication [15, 23, 24, 47]. Such distinct properties are consistent with the predictions of the model in the strong interference regime, which favors persistence of defense systems that can target competing MGEs without damaging the host.

Besides the strength of interference, a secondary factor that modulates the balance between MGE-encoded and chromosome-encoded defense systems is the scaling of the cost of defense with the number of distinct targets. Specifically, defense systems for which the cost scaling is sublinear (that is, expanding the immune repertoire is decreasingly costly) are more likely found in the chromosome. From that perspective, the relatively low prevalence of CRISPR-Cas systems in MGEs, compared to other defense systems, could be related to their capacity to expand the immune repertoire at a low cost, by simply adding spacers to the CRISPR array.

Ultra-costly defense in the strong interference regime behaves as a selfish trait that is detrimental to the host. From a global perspective, ultra costly-defense systems are also detrimental to the MGEs because, although they may increase the local abundance of their carriers, they generally decrease their overall production rate due to the damage to the host. However, from a different perspective, thanks to their ability to host ultra-costly defense systems, strongly interfering MGEs, such as incompatible plasmids, could serve as a proving ground for new defense systems. Emergent cost-inefficient defense systems could persist in these MGEs, ‘buying time’ for evolution to produce optimized (cost-efficient) variants that could then be transferred to the chromosome or other MGEs.

Although an extension of the model to more than 2 MGEs is conceptually straightforward, the enumeration of all possible combinations becomes prohibitive (there are already 81 combinations with 3 MGEs). Nevertheless, we expect that, as the number of MGEs increases, the region of the parameter space associated with strong cross-MGE interference will become broader. Indeed, the number of MGEs that a host can harbor is limited, and therefore, the more MGE classes, the more likely it is that not all of them can stably persist in coinfection. Accordingly, we would expect a higher prevalence of cost-ineffective MGE-encoded defense systems (with detrimental effects to the host), supporting complex dynamics and high MGE diversity, and facilitating the emergence of novel defense systems.

A limitation of the present work is that defense systems inducing host dormancy or regulated cell death were not considered even though such systems are common in prokaryotes [48-51]. Previous analyses have shown that, for regulated cell death to be selected as a host-level defense system, population structure or other mechanisms driving correlated sorting are required [52, 53]. We expect that similar conditions apply to the evolution of regulated cell death systems in MGEs. Thus, besides general cost-efficiency conditions, regulated cell death would only evolve in MGEs whose prevalence per ‘host patch’ (subpopulation, niche, spatial cluster) is sufficiently high.

For simplicity, the model presented here deals with a single isolated host population. More complex models could include a metapopulation structure with local co-evolution of defense and MGEs, and migration of MGEs between populations. Correlated flux of MGEs across populations might make MGE-encoded defense more efficient than chromosome-encoded defense, as it would be better (pre)adapted to changing MGE profiles. Phenomenologically, that would not be much different from having access to a lower-cost MGE-encoded defense: better adapted systems mean that less investment is needed to obtain the same overall efficacy. Nevertheless, more explicit models are required to investigate the magnitude of these effects and how they depend on cross-population fluxes and MGE diversity.

## Methods

### Numerical simulations

The model was numerically integrated with the ode45 solver of Matlab R2024b, which is based on an explicit Runge-Kutta (4,5) formula [54, 55], in a server with an Intel Xeon Gold 6248R processor. Numerical integration started from a nearly balanced initial condition, with equal fractions of H, A, B, H_a_, H_b_, H_ab_, A_b_, and B_a_ plus a small perturbation (implemented by adding a uniform random number in the interval [0,0.01] to each nonzero class and renormalizing). A baseline scenario without MGE-encoded defense was simulated by initializing the system with nearly equal fractions of H, A, B, H_a_, H_b_, and H_ab_, while setting A_b_ = B_a_ = 0. For all numerical explorations, we set *b* = 1 without loss of generality. This is equivalent to normalizing all parameters (including time) with respect to the infection rate *b*. The model was simulated for 20,000 time units. Cyclic asymptotic dynamics were identified by computing the fast Fourier transform of the last 1,000 time units and selecting the peak that corresponded to the longest period. The average population composition was then obtained using sliding windows, with window size equal to the longest period.

### Fitness calculation

A system of replicator equations can be interpreted as a model for the composition of a growing population, with the population growth rate equal to the mean fitness, Φ = Σ_*j*_ *x*_*j*_*F*_*j*_ + *M*_*j*_. Accordingly, given two conditions *a* and *b* (typically presence and absence of MGE-encoded defense), we obtained the difference in the overall growth rate as Φ^*α*^ − Φ^*β*^. Defining the fitness for individual classes and groups is more problematic because, by design of the model, they all grow at the same rate in the stationary state. Let us consider a group of classes *X* ∈ 𝒲 (for example, all hosts carrying MGE A, with and without defense systems, alone or in coinfection with B). In the stationary state, the group grows exponentially as *N*_𝒲_ (*t*) = Σ_*X*∈𝒲_ *X e*^Φ*t*^. Now let us consider two conditions, *a* and *b*, and take the log ratio 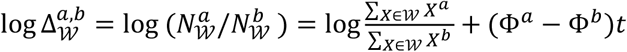 . The dependency of the last term of this expression on time underscores the difference between local (short-term) and absolute (long-term) fitness in replicator models. From a local (within-population) perspective, differences in the success of a group can be quantified by computing the log ratio of its relative abundance. From an absolute perspective, the growth rate of a group is determined by the overall growth rate of the whole population. In the case of the host, the first term always cancels out and the fitness difference only depends on Φ^*a*^ − Φ^*b*^.

## Author Contributions

Conceptualization: J.I., Y.I.W., E.V.K; model development: J.I.; investigation: J.I., Y.I.W., E.V.K.; writing – original draft: J.I; writing – review and editing: J.I., Y.I.W., E.V.K.; project administration, funding acquisition: J.I., E.V.K.

## Acknowledgements

This work was funded by MICIU/AEI/10.13039/501100011033 and by ERDF/EU (PGE) “A way of making Europe”, through grants PID2019-106618GA-I00 and CNS2023-145430 to J.I. E.V.K. and Y.I.W. are supported through the intramural program of the U.S. National Institutes of Health. The contributions of the NIH author(s) are considered Works of the United States Government. The findings and conclusions presented in this paper are those of the author(s) and do not necessarily reflect the views of the NIH or the U.S. Department of Health and Human Services.

## Appendix A From population dynamics to generalized replicator-mutator equations

We start from equation 1.

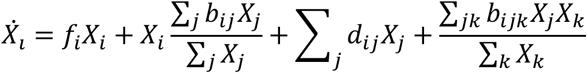

Let us define the relative abundances *x*_*i*_ = *X*_*i*_/ ∑ *X* and apply the derivative:

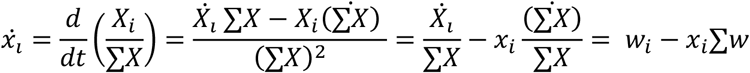

where we defined

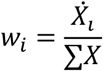

We obtain a generalized replicator-mutator equation by replacing the expression of 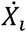 in *w*_*i*_:

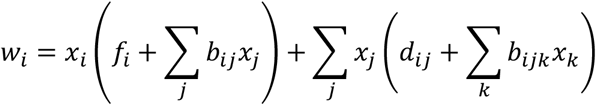

## Appendix B Explicit expressions for the generalized replicator-mutation equations

### Host death scenario

The equations that describe the dynamics of the model can be generally expressed as

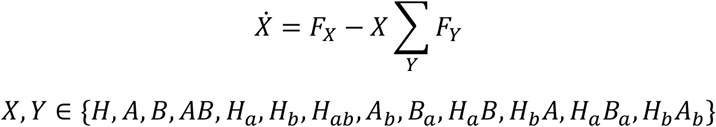

where X and Y represent each of the 13 classes of replicators and F_X_ represents the total rate of change of class X before normalization.

Assuming that the host dies if A (respectively B) is superinfected by B_a_ (respectively A_b_), the expressions for the F_X_ rates are:

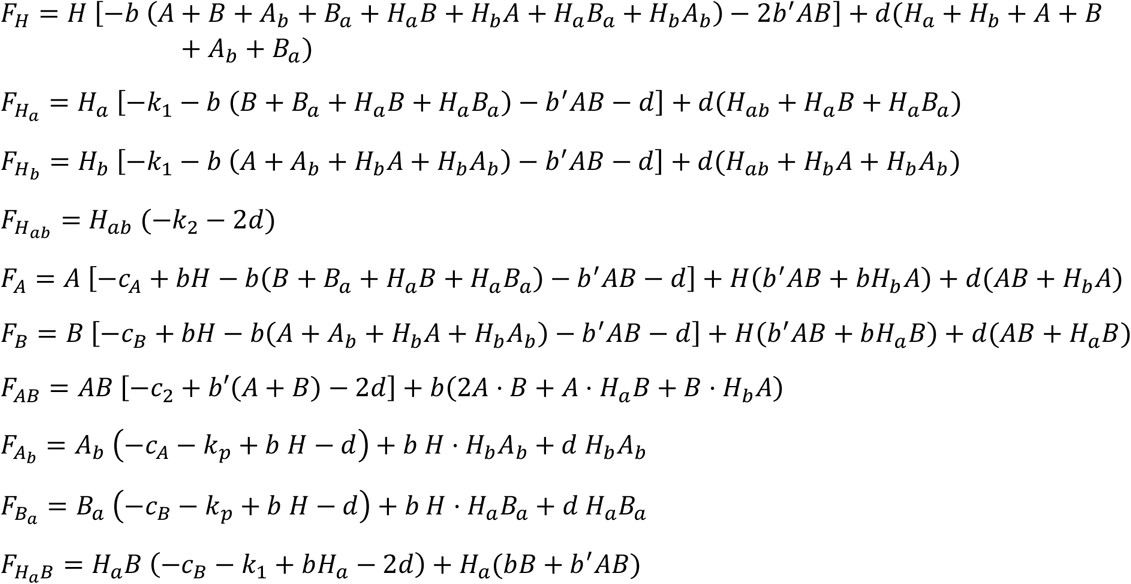

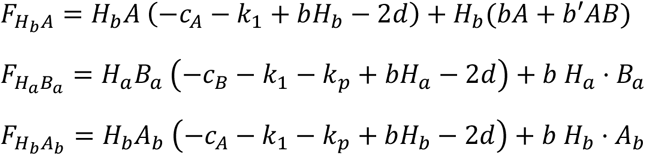

### Defense-driven replacement scenario

This scenario assumes the effective elimination of any preexisting MGE targeted by newly acquired defense systems. The same equations apply, except for 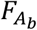 and 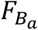:

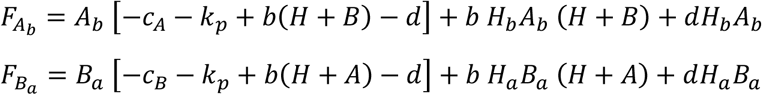

## Appendix C Coexistence of MGE in the absence of defense systems

### Stability of an MGE-free population

Let us consider an MGE-free population (entirely composed of class H) and analyze the conditions under which it cannot be invaded. To that end, we will calculate the per capita growth rate of an infinitesimal fraction of MGEs of class A, which has the most favorable transfer-to-cost ratio. Let 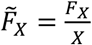 be the per capita growth rate of class X (the explicit expressions of *F*_*X*_ are provided in Appendix B). By taking the limit *H* → 1 in the generalized replicator-mutator equations we get that 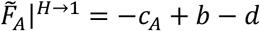 and 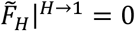. The MGE-free population remains free of A if 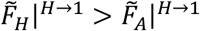, which holds if 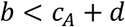.

Conversely, let us consider a pure population of class A. The per capita growth rate of an infinitesimal amount of MGE-free hosts becomes 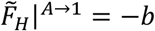 whereas the growth rate of A is 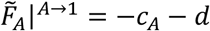. (In the expression of 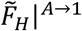 we omitted the effect of deletions, which would produce a residual population of non-self-sustaining MGE-free hosts.) Comparing both growth rates, it comes out that MGE-free hosts can invade the population if *b* < *c*_*A*_ + *d*.

Combining both results, and as long as *b*^′^ ≤ *b, c*_*B*_ ≥ *c*_*A*_, and *c*_2_ ≥ *c*_*A*_ (as reasonably expected), the condition *b* < *c*_*A*_ + *d* implies that the population will be entirely composed of MGE-free hosts.

### Fixation of a single MGE

Following the same approach, we can study the stability of a population of hosts that carry MGE class A. Based on the expressions obtained above, the condition for a small fraction of A to invade an MGE-free population is *b* > *c*_*A*_ + *d*. The same condition must be held so that a population entirely composed of class A is not invaded by class H.

On the other side, class A can invade a population entirely composed of class AB if 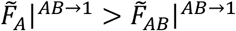, and resist invasion by that class if 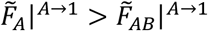, where 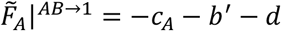, 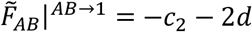, and 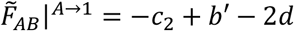 (as before, we omitted deletion terms that would lead to residual non-self-sustaining subpopulations). By combining these expressions, we conclude that class A outcompetes AB if *b*^′^ < *c*_2_ − *c*_*A*_ + *d*.

Finally, let us consider the competition between A and B. Because 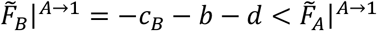, a population of class A is always resistant to invasion by B. Conversely, a small fraction of A can invade a population of class B if 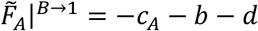, where 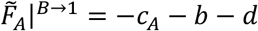 and 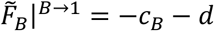. This latter condition can be expressed as *c*_*B*_ − *c*_*A*_ > *b*. If that condition does not hold, neither A can invade B nor B can invade A. In particular, A cannot invade B if *b* > *c*_*B*_ + *d* (which is the condition for B to invade an MGE-free population). As a result, for parameter combinations that simultaneously fulfil *b* > *c*_*B*_ + *d* and *b*^′^ < *c*_2_ − *c*_*B*_ + *d*, the system is bistable, with pure populations of class A and class B as alternative solutions depending on the initial composition. By solving the equation 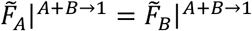 it is possible to conclude that, starting from a population that only contains classes A and B, the final composition will be all-A if 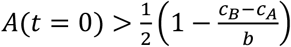and all-B otherwise. As a corollary, if the initial composition is balanced, the typical outcome will be all-A except if *c*_*A*_ ≈ *c*_*B*_. Also, for a fixed ratio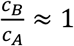and nearly balanced initial composition, the probability of reaching an all-B stationary state decreases with the average cost of the MGEs as Pr(*B*) ∝ −2*δc*_1_, where 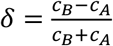.

### Stability of class AB

Finally, let us consider the class of hosts that carry both MGEs in every genome (class AB). Using the expressions derived above, we conclude that a small fraction of AB can invade a population of class A if *b*^′^ > *c*_2_ − *c*_*A*_ + *d*. (Because *c*_*A*_ ≤ *c*_*B*_ by design, the condition to invade a population of class B, that is *b*^′^ > *c*_2_ − *c*_*B*_ + *d*, is more permissive. However, if *b*^′^ > *c*_2_ − *c*_*B*_ + *d* but *b*^′^ < *c*_2_ − *c*_*A*_ + *d*, class B will be replaced by A, not by AB.) The same condition guarantees that a population of class AB cannot be invaded by A or B. Finally, AB can resist invasion by MGE-free hosts if 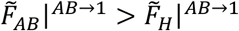, where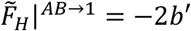The latter condition can be expressed as 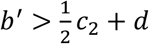 and is automatically fulfilled if *b*^′^ > *c*_2_ − *c*_*A*_ + *d*, as long as *c*_2_ ≥ 2*c*_*A*_ (which holds for all parameter combinations explored in this work). In conclusion, if *b*^′^ > *c*_2_ − *c*_*A*_ + *d*, the population will be entirely composed of hosts carrying both MGEs in every genome.

## Appendix D Analytical conditions for the spread of chromosome-encoded defense

To analyze the stability of chromosome-encoded defense, we distinguish between two qualitatively different regimes: (i) when only one MGE reaches fixation in the absence of defense, and (ii) when both MGEs are present in all genomes (fixation of class AB). As shown in Appendix C, the first regime corresponds to the conditions *b* > *c*_*A*_ + *d* and *b*^′^ < *c*_2_ − *c*_*A*_ + *d*, whereas the second regime occurs if *b*^′^ > *c*_2_ − *c*_*A*_ + *d*.

### Single-MGE regime (strong interference between MGEs)

For simplicity, we will focus on the case in which MGEs of class A reach fixation (the procedure for class B is analogous). A necessary condition for the spread of chromosome-encoded defense in this regime is that the per capita growth rate of hosts carrying a defense system against A is greater than the per capita growth rate of class A. This condition can be expressed as 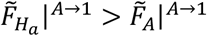, where 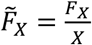 and the limit *A* → 1 represents a pure-A population. By replacing the values of *F*_*X*_ from Appendix B, we obtain the condition *k*_1_ < *c*_*A*_. This expression admits a simple interpretation in terms of cost-benefit balance: for chromosome-encoded immunity to spread, its cost must be lower than the cost of the MGE being targeted.

### Double-MGE regime (weak interference between MGEs)

Chromosome-encoded defense in this regime can appear in two ways: (i) as double immunity against both MGEs (class H_ab_) or (ii) as single immunity against one MGE while still carrying the other MGE (classes H_b_A or H_a_B). In the case of double immunity, a necessary condition for the spread of defense is 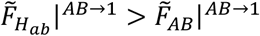, which results in *k*_2_ < *c*_2_. In the case of single immunity, the condition becomes 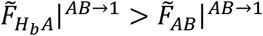, leading to *k*_1_ < *c*_2_ − *c*_*A*_. (An analogous calculation for the spread of class H_a_B leads to *k*_1_ < *c*_2_ − *c*_*B*_, which is more restrictive than the condition for H_b_A.) As in the single-MGE regime, the conditions *k*_2_ < *c*_2_ and *k*_1_ < *c*_2_ − *c*_*A*_ have a straightforward interpretation as cost-benefit balances, such that the cost of defense must be lower than the cost of the MGEs being targeted.

## Appendix E Analytical conditions for the spread of MGE-encoded defense

In this section, we derive necessary conditions for the spread of MGE-encoded defense systems in the absence of chromosome-encoded defense. First, we analyze the weak interference regime, in which both MGEs reach fixation in all genomes (in the absence of defense). Then, we analyze the strong interference regime, in which only one MGE reaches fixation in the absence of defense.

### Weak interference, no chromosome-encoded defense

As shown in Appendix C, if *b*^′^ > *c*_2_ − *c*_*A*_ + *d* both MGEs will invade all genomes, leading to a pure AB population. In this regime, chromosome-encoded defense against both MGEs (class H_ab_) is stable if *k*_2_ < *c*_2_, whereas chromosome-encoded defense against a single MGE (class H_b_A) is stable if *k*_1_ < *c*_2_ − *c*_*A*_. Hosts belonging to any of these classes (AB, H_ab_, H_b_A) are resistant to further infection due to superinfection exclusion, immunity, or a combination of both. Therefore, MGE-encoded defense can only spread if the per capita growth rate of defense-encoding MGEs (classes A_b_ and B_a_) is higher than the per capita growth rate of the resident class. In other words, in this scenario the spread of MGE-encoded defense occurs only through competition among hosts.

Let us focus on the case in which chromosome-encoded defense is absent. The per capita growth rate of a small fraction of class A_b_ in a population of class AB is 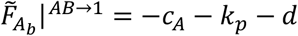, whereas the per capita growth rate of the AB population is 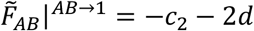. Thus, MGE-encoded defense will spread if *k*_*p*_ < *c*_2_ − *c*_*A*_ + *d*. (The condition for the spread of B_a_ is *k*_*p*_ < *c*_2_ − *c*_*B*_ + *d*, which is more restrictive.) The maximum possible cost of MGE-encoded defense becomes 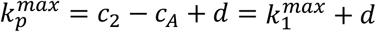. In this scenario MGE-encoded defense will not evolve if it is more costly than chromosome-encoded defense plus the loss rate. The resemblance between the expressions found for chromosome-encoded defense and MGE-encoded defense underscores the fact that inter-MGE competition in this regime only occurs at the host level. In both cases, immunity (regardless of whether it is encoded in the chromosome or an MGE) pays off if it is less costly than the MGE that is being targeted. After some manipulation, the expression of 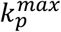can be rewritten as 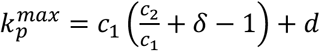, where 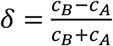. Thus, for fixed 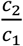 and 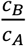, the maximum cost of MGE-encoded defense becomes proportional to the average cost of an MGE.

### Strong interference, no chromosome-encoded defense

If *b*^′^ < *c*_2_ − *c*_*A*_ + *d* only one MGE reaches fixation in the population (Appendix C). Unlike in the weak interference regime, the single-MGE steady state can be destabilized by two mechanisms. On one hand, hosts carrying a single MGE can be outcompeted by other classes of faster-growing hosts (host-driven competition). On the other hand, hosts carrying a single MGE can be further infected by the other MGE, modifying the intra-host MGE composition (infection-driven competition). This second level of competition is critically dependent on the superinfection scenario.

Let us first analyze the host death scenario. If we assume an all-A population, then we have that 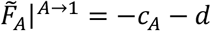 and 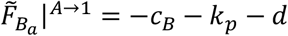. Therefore, a necessary condition for the spread of B_a_ is *k*_*p*_ < *c*_*A*_ − *c*_*B*_. By design, *c*_*A*_ < *c*_*B*_, which implies that MGE-encoded defense cannot spread in this case. In contrast, an all-B population allows the spread of A_b_ if *k*_*p*_ < *c*_*B*_ − *c*_*A*_. As discussed in Appendix C, the all-B scenario is only relevant if *c*_*A*_ ≈ *c*_*B*_ or if the initial composition is sufficiently skewed towards B.

In the target replacement scenario, the per capita growth rate of B_a_ has an additional term associated with infection and replacement of class A, such that 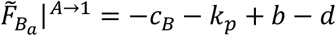. The condition for B_a_ to spread becomes *k*_*p*_ < *b* + *c*_*A*_ − *c*_*B*_. (In the case of an all-B population, the condition for A_b_ to spread would be *k*_*p*_ < *b* + *c*_*B*_ − *c*_*A*_.) After some manipulation, the maximum cost of MGE-encoded defense can be expressed as 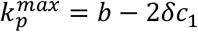, where 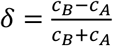. In this scenario, 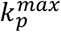 decreases with the average cost of MGEs, although such decrease is negligible if *c*_*A*_ ≈ *c*_*B*_. Note that the infection term *b* is the ultimate reason why 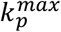 can reach values substantially higher than *k*_1_ in this scenario.

### Parameter regions compatible with chromosome-encoded defense

If there is chromosome-encoded defense, the situation becomes more complicated due to the establishment of cyclic dynamics involving fluctuating fractions of single-MGE classes. A comprehensive analytical study of these cases based on invasibility arguments becomes unfeasible, as it would require knowing the nature of the cycles generated by each parameter combination and, even if so, taking the limit *X* → 1 in the calculation of 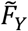 would not be justified in general. Nevertheless, the presence of single-MGE classes in cyclic regimes implies that the spread of MGE-encoded defense has an infection component, which promotes the spread of MGE-encoded defense in the target replacement scenario. In consequence, we expect to observe major differences between alternative superinfection scenarios, with higher values of 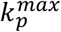 (and possibly 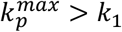) in the case of target replacement.

In some special cases, it is possible to obtain analytical expressions of 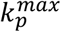 in regions in which chromosome-encoded defense could evolve. The key is to identify parameter regions in which the per capita growth rates of the classes of interest do not depend on the quantitative composition of the population. That occurs, for example, in the weak interference regime (*b*^′^ > *c*_2_ − *c*_*A*_ + *d*) if the additional condition *k*_1_ > *c*_*A*_ holds, which guarantees the absence of classes H_a_ and H_b_ (Appendix D). Let us denote that region of the parameter space as Γ. Simulations show that, in this region, class H is also absent. The presence of class H_b_A_b_ can be discarded by applying the same arguments developed below. The per capita growth rates of classes H_ab_ and H_b_A become 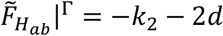and 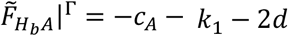. In the host death scenario, the per capita growth rate of class A_b_ is 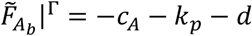. The fact that these three growth rates are independent of the quantitative composition of the population implies that H_ab_, H_b_A, and A_b_ cannot coexist. The defense-encoding MGE class A_b_ will outcompete H_ab_ and H_b_A if *k*_*p*_ < min{*k*_2_ − *c*_*A*_ + *d*; *k*_1_ + *d*}. The value of 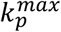 obtained in the absence of chromosome-encoded defense is smaller than this expression if there is no defense and greater than it if there is defense. As a result, the maximum cost of MGE-encoded defense in the parameter region Γ under the host death scenario can be written as 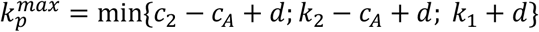. A similar derivation is not possible in the target replacement scenario because, in that case, the per-capita growth rate of class A_b_ depends on the abundance of class B 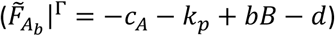, which cannot be easily determined a priori. The expression above still provides a lower bound for 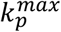, whereas an upper bound would be given by 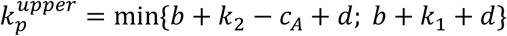 in the region in which chromosome encoded defense can evolve (defined by *k*_2_ < *c*_2_ or *k*_1_ < *c*_2_ − *c*_*A*_) and 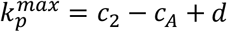otherwise.

