## Supplementary Figures for "Eco-evolutionary dynamics of defense systems in mobile genetic elements: Cui bono?"

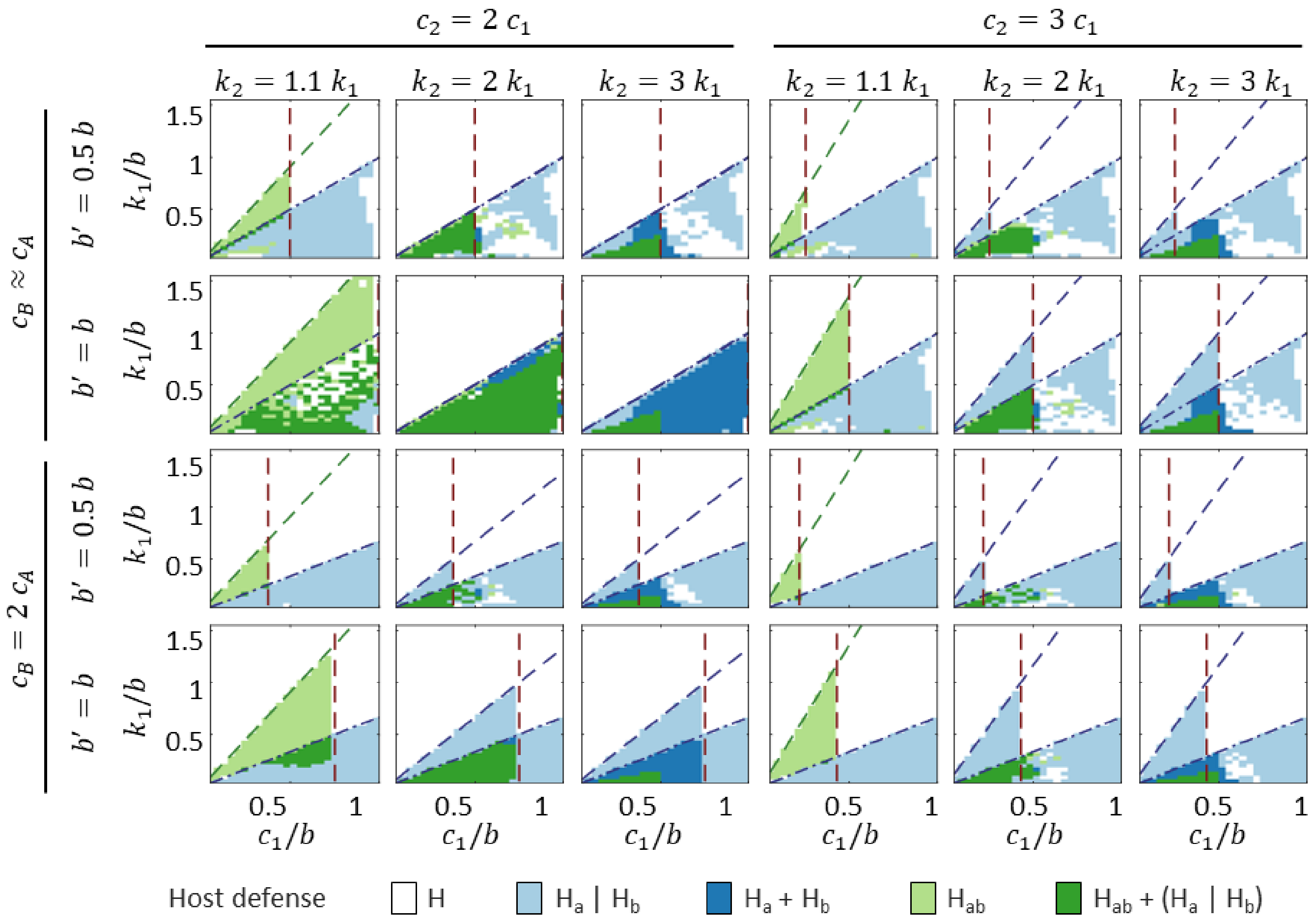


**Figure S1:** Chromosome-encoded defense in a population with 2 MGEs when MGE-encoded defense is not available. "$H_a$ | $H_b$" denotes the presence of one of those classes, but not both. Simulations were run with zero loss rate ($d = 0$). Brown dashed line: limit condition separating the weak (left) and strong (right) interference regimes, $R_0^{AB} = 1$. Dashed green line: limit condition for the spread of defense against both MGEs, $k_2 = c_2$. Dash-dotted blue line: limit condition for the spread of defense against a single MGE, $k_1 = c_A$. Dashed blue line: limit condition for the spread of defense against one MGE in hosts already carrying the other MGE, $k_1 = c_2 - c_A$.

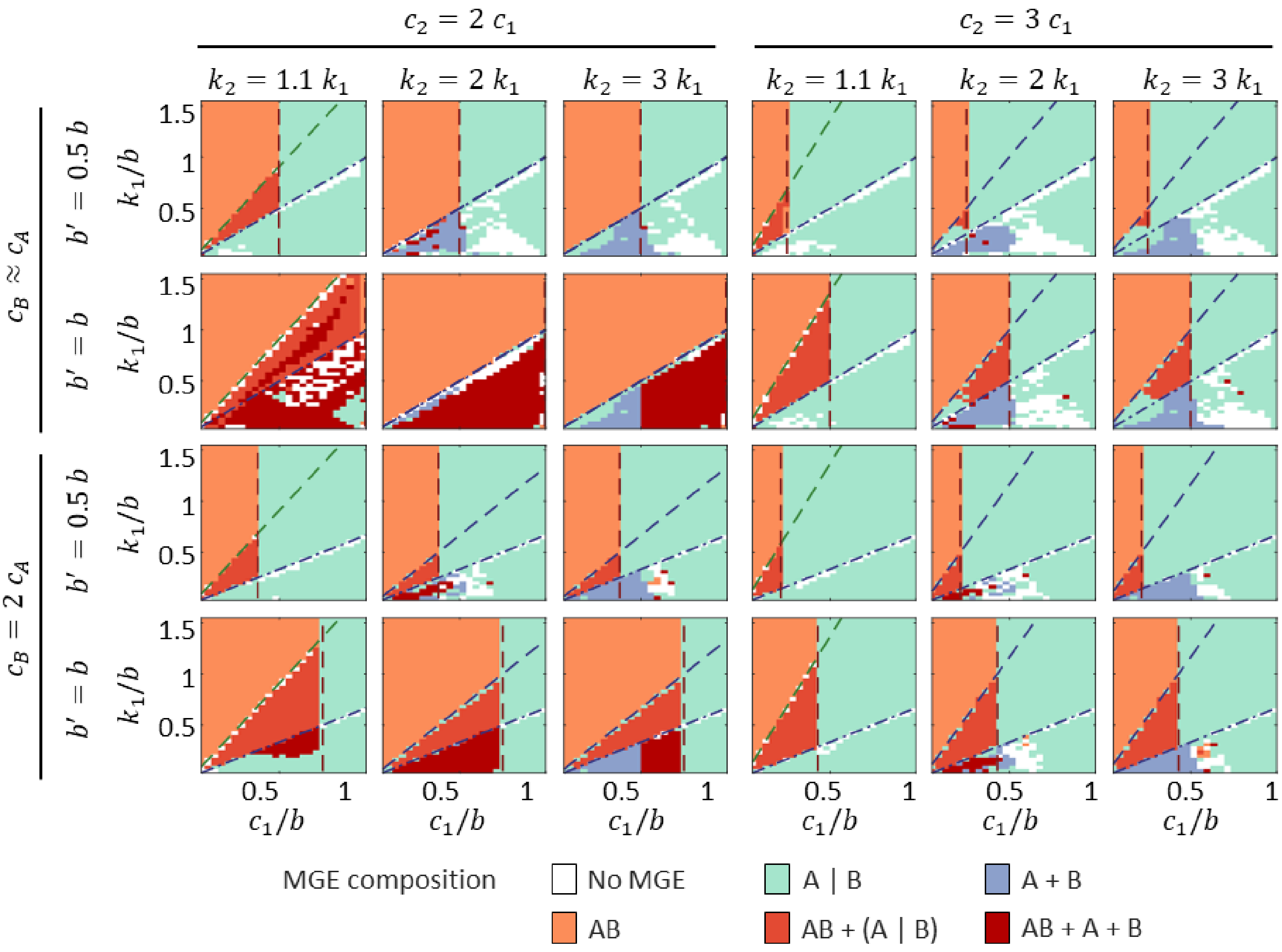


**Figure S2:** Composition of the MGE population in the absence of MGE-encoded defense (A). "A|B" denotes the presence or MGE classes A or B, but not both. Simulations were run with zero loss rate ($d = 0$). Brown dashed line: limit condition separating the weak (left) and strong (right) interference regimes, $R_0^{AB} = 1$. Dashed green line: limit condition for the spread of defense against both MGEs, $k_2 = c_2$. Dash-dotted blue line: limit condition for the spread of defense against a single MGE, $k_1 = c_A$. Dashed blue line: limit condition for the spread of defense against one MGE in hosts already carrying the other MGE, $k_1 = c_2 - c_A$.

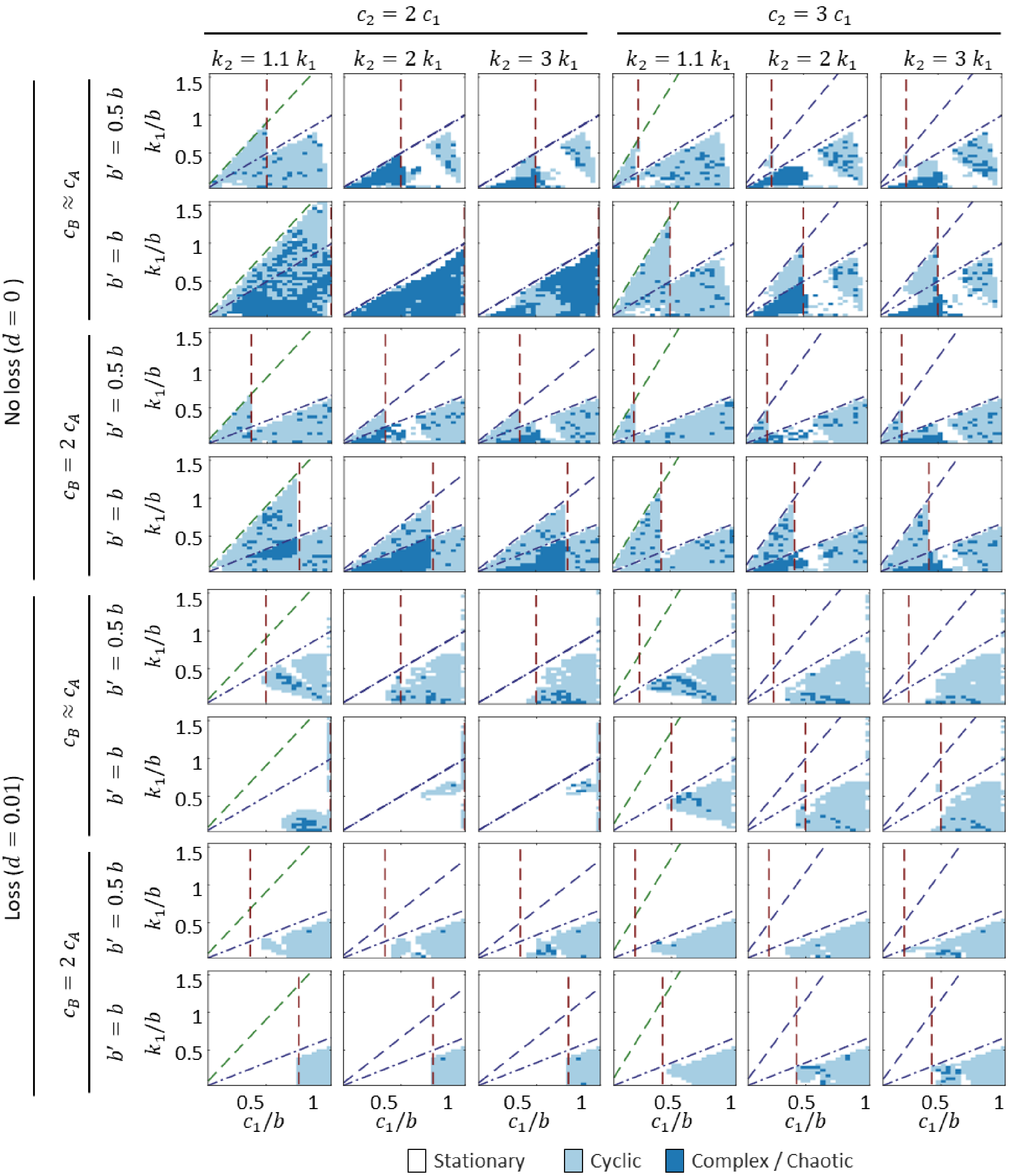


**Figure S3:** Periodicity of the MGE population dynamics in the absence of MGE-encoded defense, with and without gene loss. Brown dashed line: limit condition separating the weak (left) and strong (right) interference regimes, $R_0^{AB} = 1$. Dashed green line: limit condition for the spread of defense against both MGEs, $k_2 = c_2$. Dash-dotted blue line: limit condition for the spread of defense against a single MGE, $k_1 = c_A$. Dashed blue line: limit condition for the spread of defense against one MGE in hosts already carrying the other MGE, $k_1 = c_2 - c_A$.

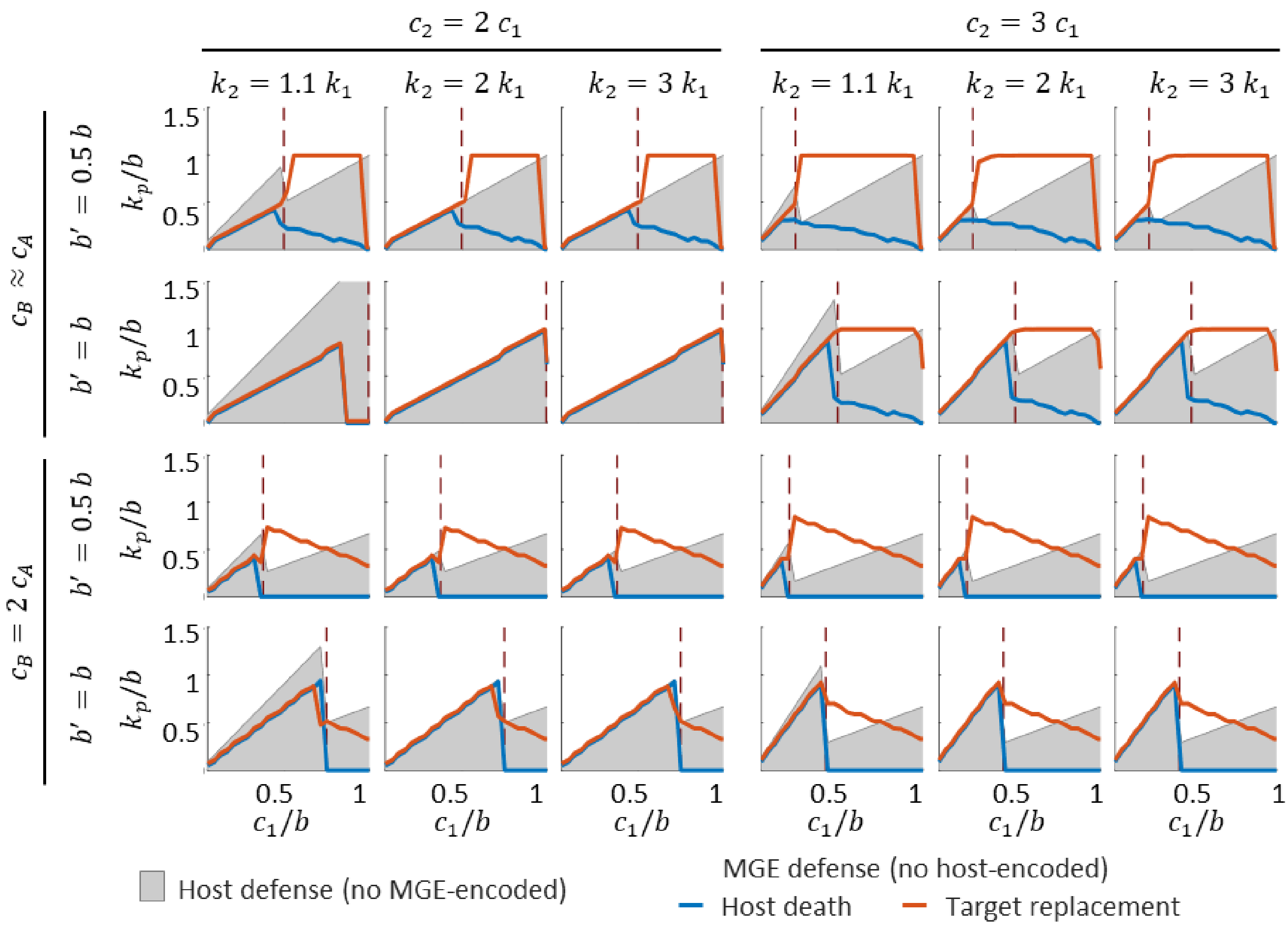


**Figure S4:** Maximum possible cost of MGE-encoded defense in the absence of chromosome-encoded defense. Simulations were run with zero loss rate ($d = 0$). For comparison, regions in which chromosome-encoded defense would be cost-efficient are shown in gray. Brown dashed line: limit condition separating the weak (left) and strong (right) interference regimes, $R_0^{AB} = 1$.

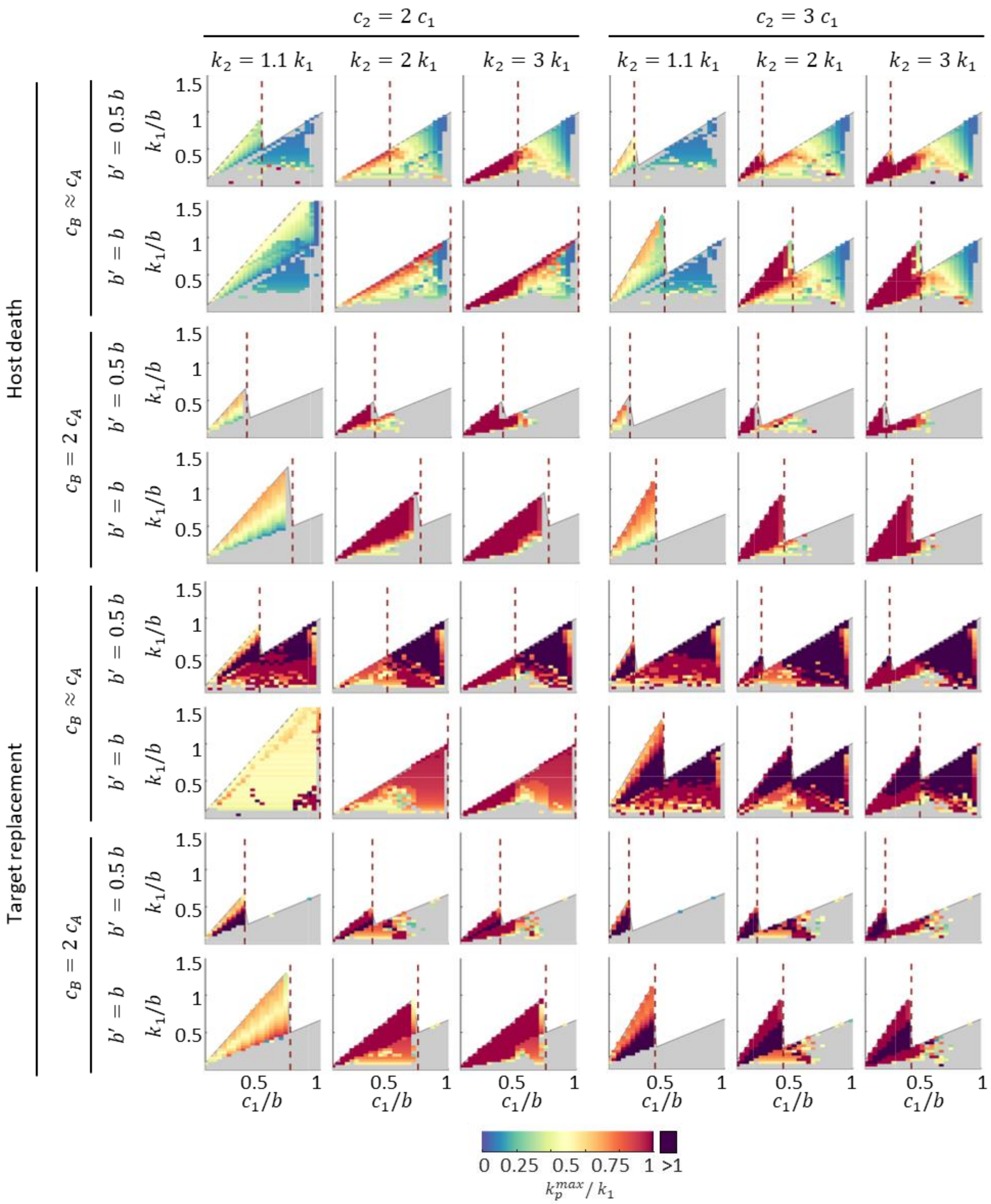


**Figure S5:** Maximum possible cost of MGE-encoded defense ($k_p^{max}$) relative to the cost of chromosome-encoded defense. Simulations were run with zero loss rate ($d = 0$). Brown dashed line: limit condition separating the weak (left) and strong (right) interference regimes, $R_0^{AB} = 1$. Regions in gray correspond to parameter combinations in which chromosome-encoded defense is cost-efficient and would evolve in the absence of MGE-encoded defense.

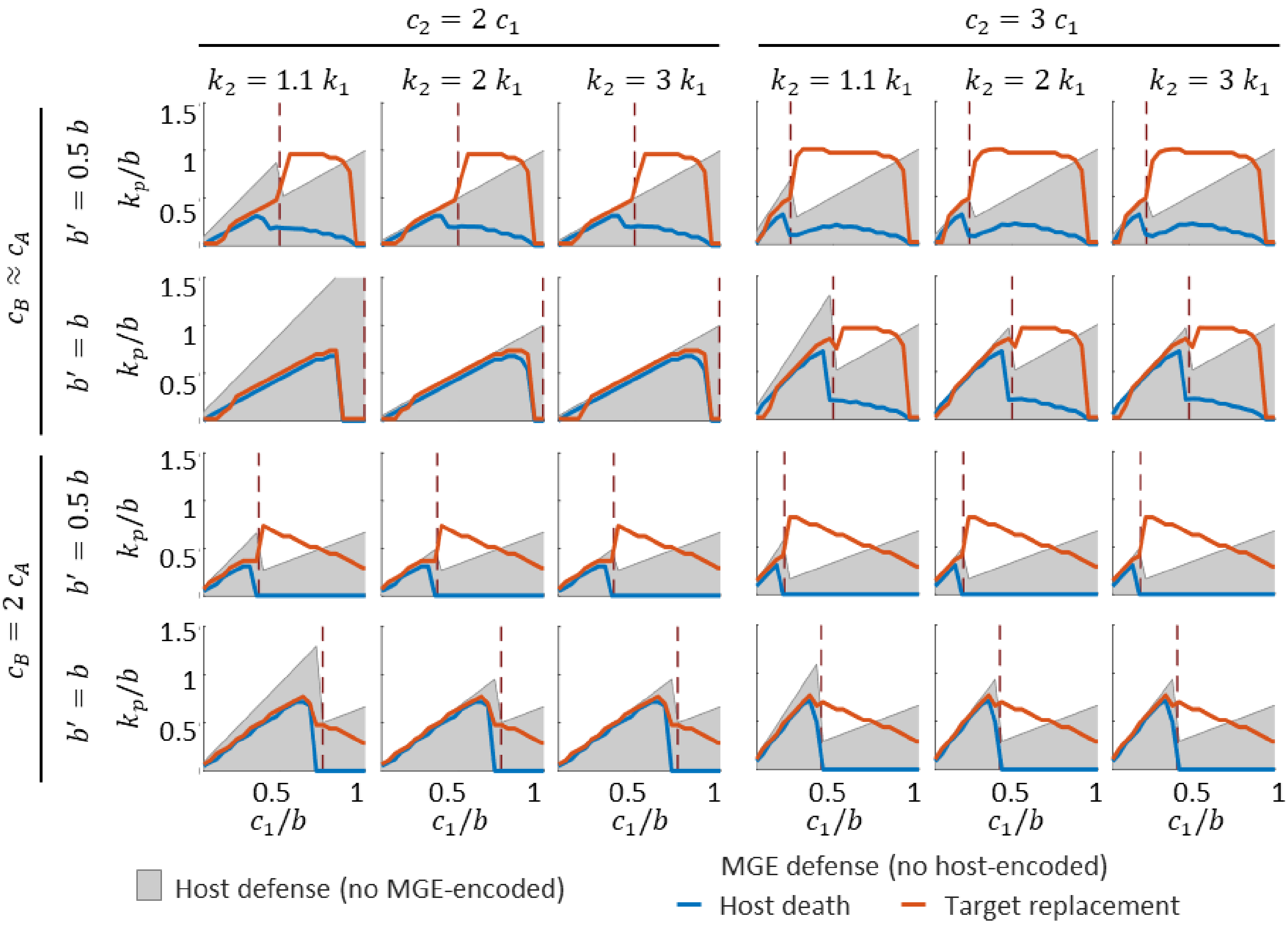


**Figure S6:** Maximum possible cost of MGE-encoded defense in the absence of chromosome-encoded defense. Simulations were run with nonzero deletion rate ($d = 0.01$). For comparison, regions in which chromosome-encoded defense would be cost-efficient are shown in gray. Brown dashed line: limit condition separating the weak (left) and strong (right) interference regimes, $R_0^{AB} = 1$.

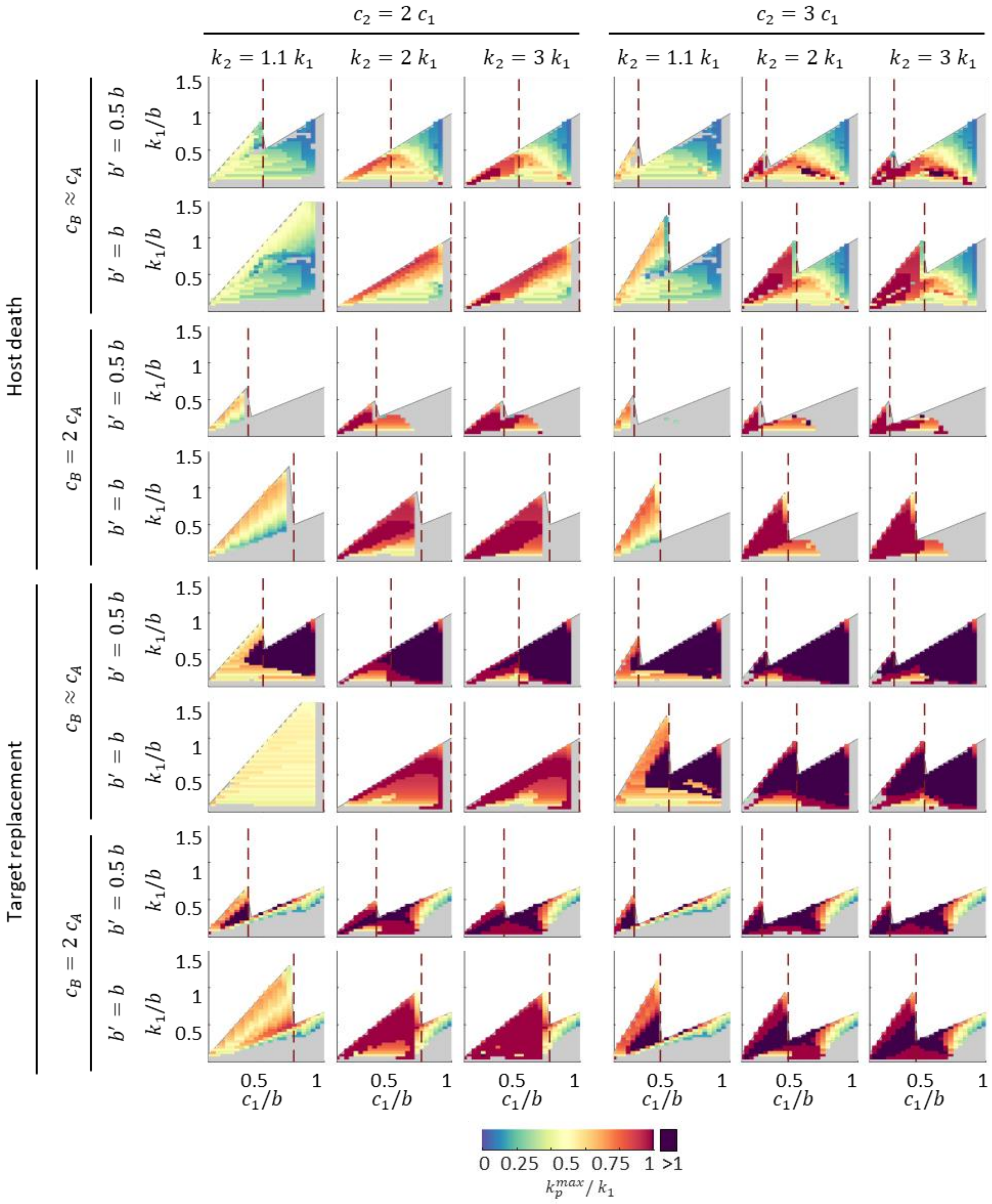


**Figure S7:** Maximum possible cost of MGE-encoded defense ($k_p^{max}$) relative to the cost of chromosome-encoded defense. Simulations were run with nonzero deletion rate ($d = 0.01$). Brown dashed line: limit condition separating the weak (left) and strong (right) interference regimes, $R_0^{AB} = 1$. Regions in gray correspond to parameter combinations in which chromosome-encoded defense is cost-efficient and would evolve in the absence of MGE-encoded defense.

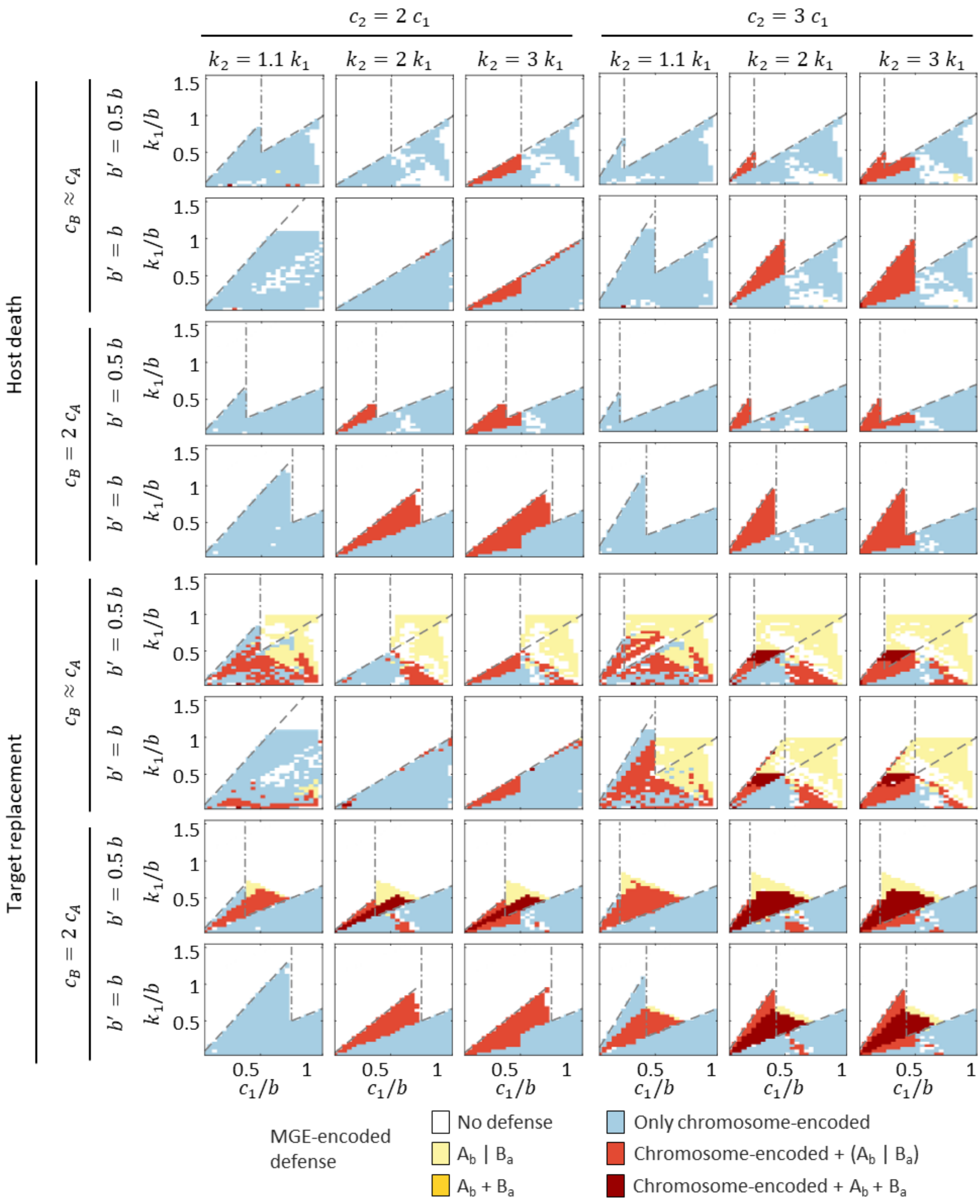


**Figure S8:** Expected location of defense in the long-term, considering the possible transfer of MGE-encoded defense to the chromosome and vice versa. "$A_b$ | $B_a$" indicates that only one of the two MGEs will encode a defense system. Brown dashed line: limit condition separating the weak (left) and strong (right) interference regimes, $R_0^{AB} = 1$. Dashed lines separate regions in which chromosome-encoded defense is cost-efficient or not (below and above the line, respectively). All simulations were run with zero loss rate ($d = 0$) and the same cost for MGE-encoded defense and chromosome-encoded defense ($k_p = k_1$).

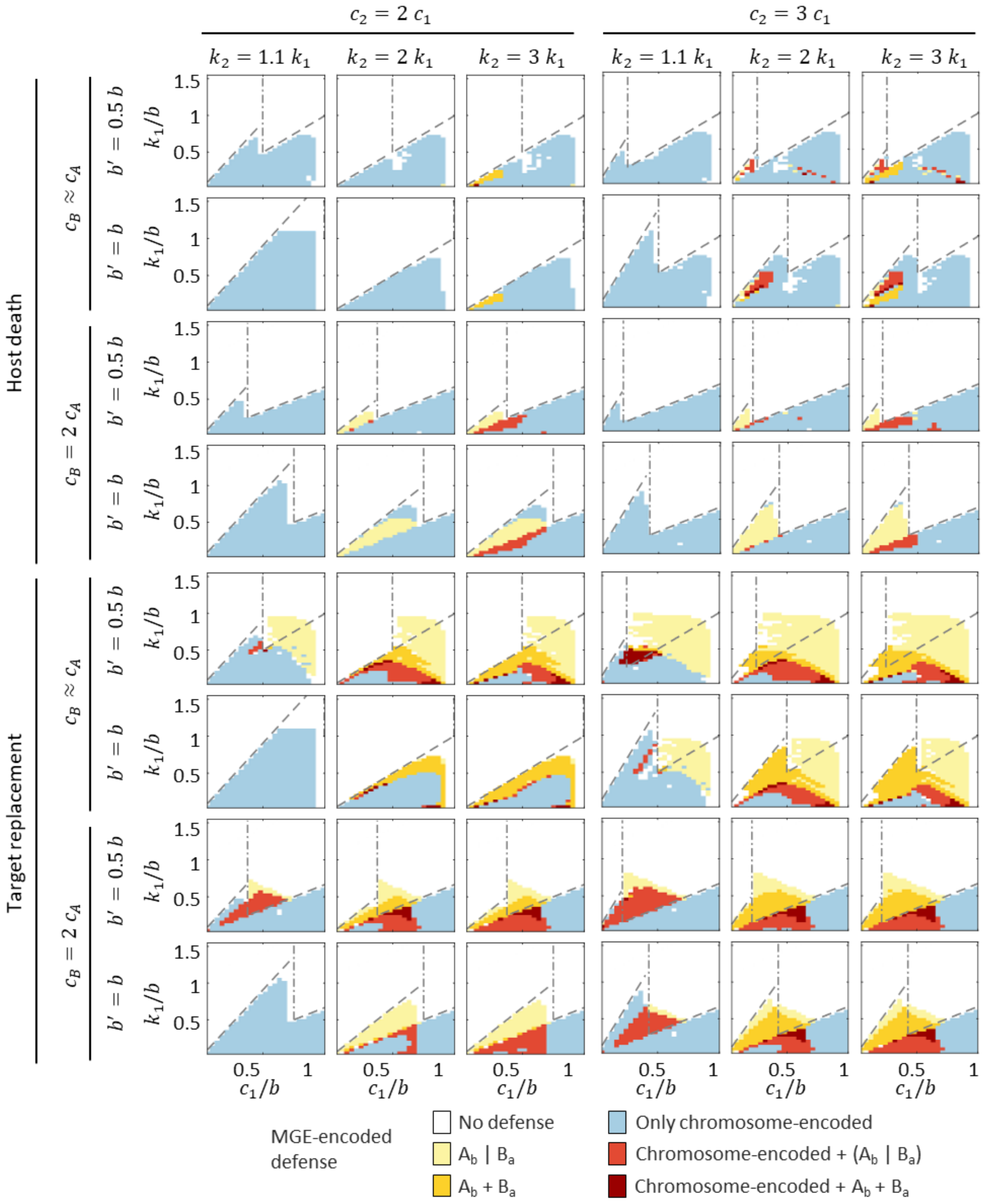


**Figure S9:** Expected location of defense in the long-term, considering the possible transfer of MGE-encoded defense to the chromosome and vice versa. "$A_b \mid B_a$" indicates that only one of the two MGEs will encode a defense system. Brown dashed line: limit condition separating the weak (left) and strong (right) interference regimes, $R_0^{AB} = 1$. Dashed lines separate regions in which chromosome-encoded defense is cost-efficient or not (below and above the line, respectively). All simulations were run with nonzero loss rate ($d = 0.01$) and the same cost for MGE-encoded defense and chromosome-encoded defense ($k_p = k_1$).

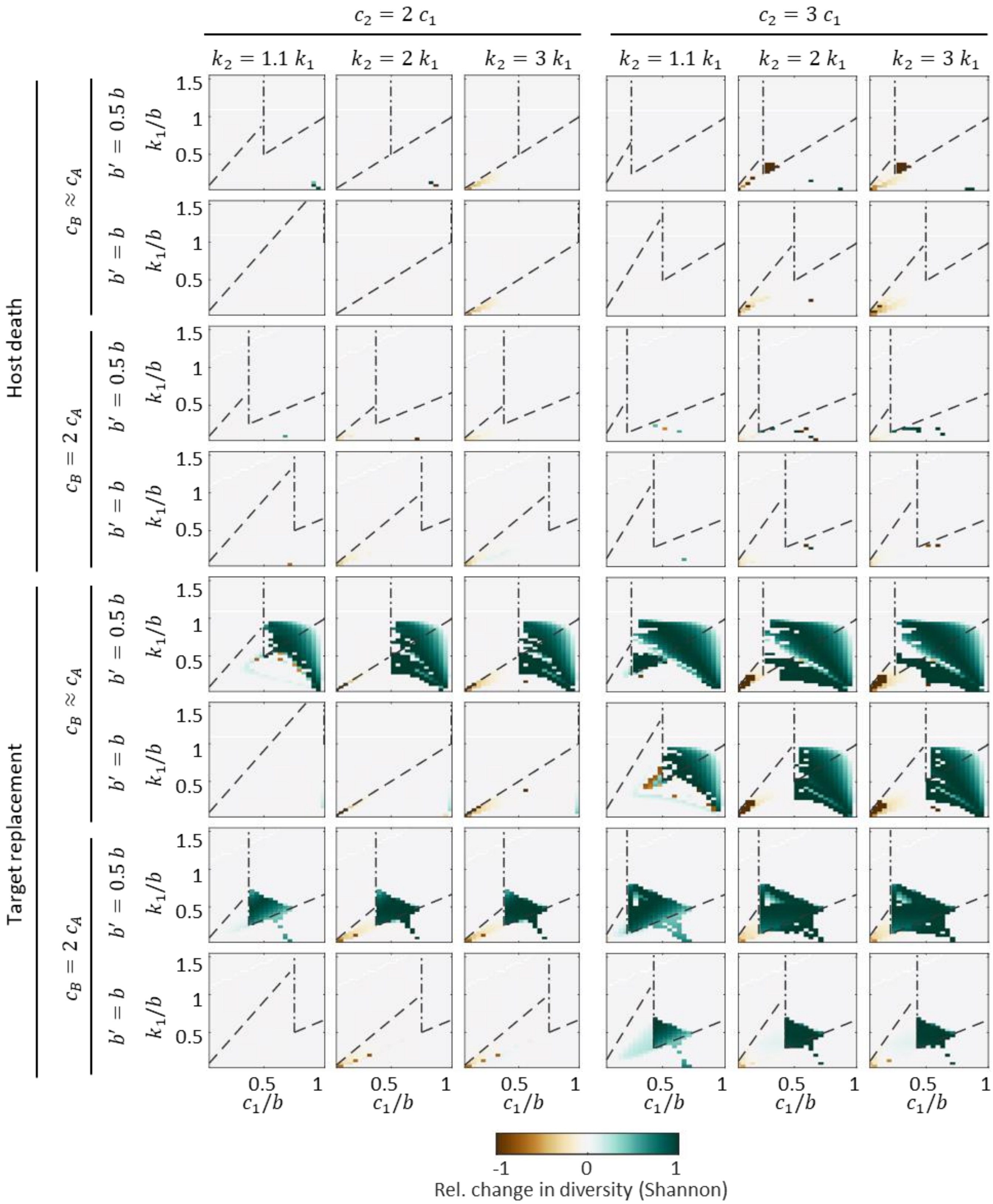


**Figure S10:** Effect of MGE-encoded defense on the population-level diversity of MGEs. Diversity was quantified using the Shannon index, jointly considering all variants (defense-free and defense-encoding) of each class of MGE. Simulations were run with nonzero deletion rate ($d = 0.01$) and same cost for MGE-encoded defense and chromosome-encoded defense ($k_p = k_1$). Vertical dash-dotted line: limit condition separating the weak (left) and strong (right) interference regimes, $R_0^{AB} = 1$. Dashed lines separate regions in which chromosome-encoded defense is cost-effective or not (below and above the line, respectively).

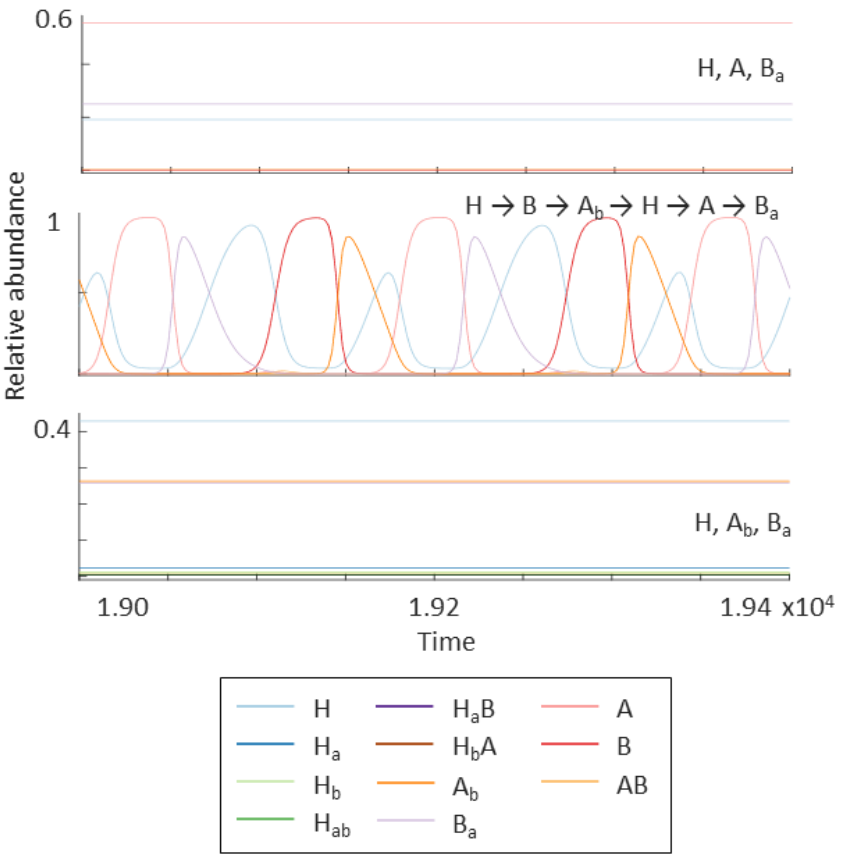


**Figure S11:** Representative examples of MGE coexistence driven by MGE-encoded defense. Parameter values: $c_1 = 0.7$, $k_1 = 0.8$ (top), $k_1 = 0.4$ (middle), $k_1 = 0.25$ (bottom), $k_p = k_1$, $b = 1$, $b' = 0.5$, $c_2 = 2c_1$, $k_2 = 2k_1$, $c_B = 1.01\ c_A$, $d = 0.01$. All simulations correspond to the target replacement scenario.

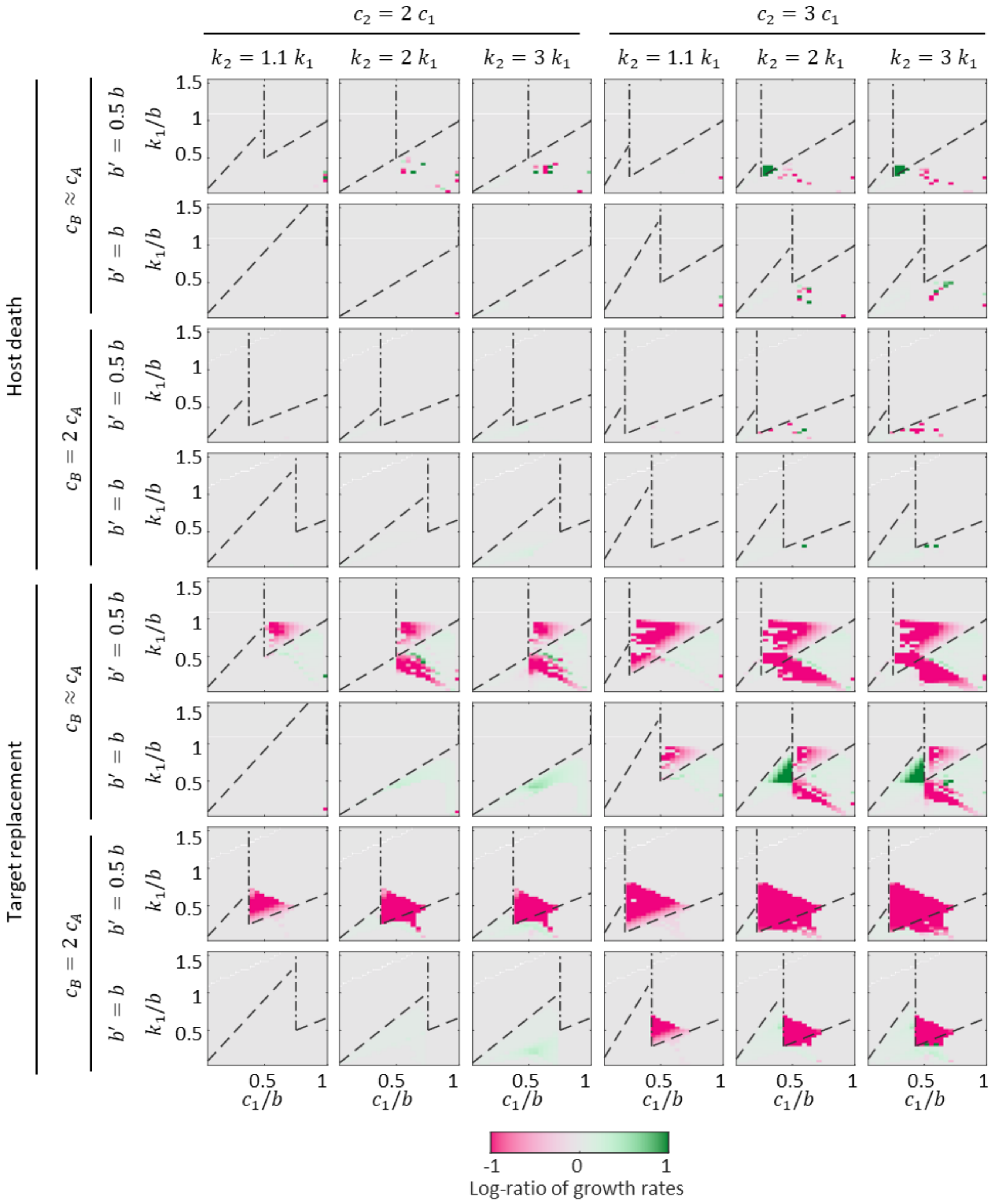


**Figure S12:** Effect of MGE-encoded defense on the overall fitness of the population, quantified as the change in the logarithm of the growth rate after introducing MGE-encoded defense. Simulations were run with nonzero deletion rate ($d = 0.01$) and same cost for MGE-encoded defense and chromosome-encoded defense ($k_p = k_1$). Vertical dash-dotted line: limit condition separating the weak (left) and strong (right) interference regimes, $R_0^{AB} = 1$. Dashed lines separate regions in which chromosome-encoded defense is cost-effective or not (below and above the line, respectively).

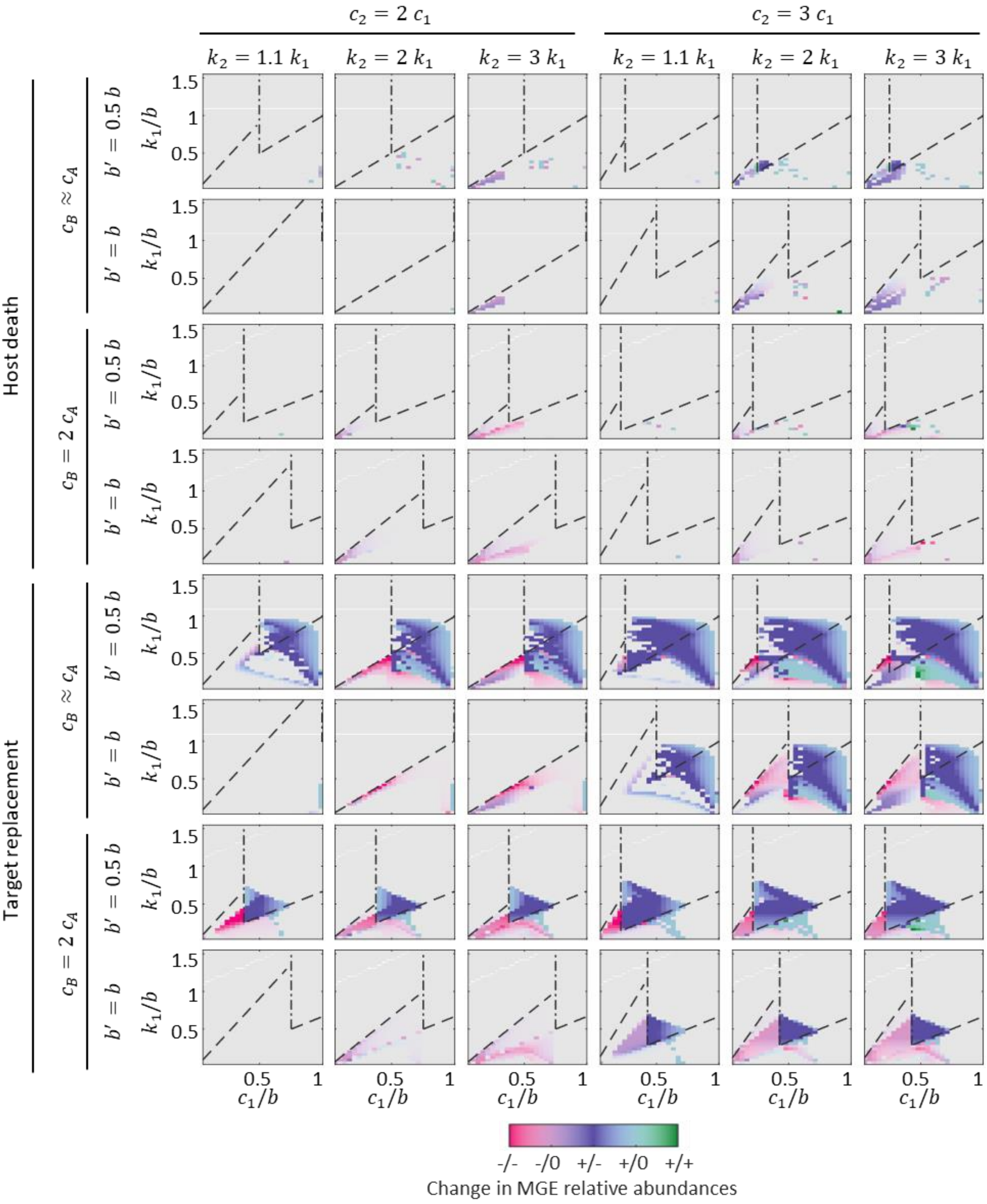


**Figure S13:** Effect of MGE-encoded defense on the local fitness of each MGE, quantified as the change in their relative abundance after introducing MGE-encoded defense. To calculate the relative abundances, all variants (defense-free and defense-encoding) of each class of MGE were jointly considered. Simulations were run with nonzero deletion rate ($d = 0.01$) and same cost for MGE-encoded defense and chromosome-encoded defense ($k_p = k_1$). Vertical dash-dotted line: limit condition separating the weak (left) and strong (right) interference regimes, $R_0^{AB} = 1$. Dashed lines separate regions in which chromosome-encoded defense is cost-effective or not (below and above the line, respectively).
